# Evolution of warfare by resource raiding favors polymorphism in belligerence and bravery

**DOI:** 10.1101/2021.11.01.466436

**Authors:** Charles Mullon, Laurent Lehmann

## Abstract

From protists to primates, intergroup aggression and warfare over resources has been observed in several taxa whose populations typically consist of groups connected by limited genetic mixing. Here, we model the co-evolution between four traits relevant to this setting: (i) investment into common-pool resource production within groups (“helping”); (ii) proclivity to raid other groups to appropriate their resources (“belligerence”); and investments into (iii) defense and (iv) offense of group contests (“defensive and offensive bravery”). We show that when traits co-evolve, the population often experiences disruptive selection favouring two morphs: “Hawks”, who express high levels of both belligerence and offensive bravery; and “Doves”, who express neither. This social polymorphism involves further among-traits associations when the fitness costs of helping and bravery interact. In particular if helping is antagonistic with both forms of bravery, co-evolution leads to the coexistence of individuals that either: (i) do not participate into common-pool resource production but only in its defense and appropriation (“Scrounger Hawks”); or (ii) only invest into common pool resource production (“Producer Doves”). Provided groups are not randomly mixed, these findings are robust to several modelling assumptions. This suggests that inter-group aggression is a potent mechanism in favoring within-group social diversity and behavioural syndromes.

## 1 Introduction

Warfare –the coalitionary aggression between groups of individuals – is one of the defining features of the human lineage. It is not only thought to have driven the advent of large-scale societies, but also that small scale hunter-gatherer societies regularly took part in coalitionary contests for material resources and reproductive opportunities [1–3]. Warfare is also well known to occur in chimpanzees [4], and has further been observed in several taxa outside of primates: in banded mongoose, where it is initiated by females who seek to mate with extragroup males [5]; in army ants, where colonies engage in contests for access to new territories [6]; and even in some strains of bacteria, who use a large assemblage of different offensive weapons to dislodge patches of rival cells [7]. As highlighted by these examples, warfare typically involves a resource over which conflict occurs between groups, and such conflict depends on individual contributions into offensive and defensive group-level activities [8]. This leads to an overall structure of interactions involving in-group solidarity and out-group hostility.

There is a large theoretical literature on the behavioral underpinnings of warfare that use different approaches to consider related questions about within-group solidarity and between-group hostility. In economics and game theory, models often focus on understanding the Nash equilibrium strategies for two types of behaviours under different scenarios of intergroup conflict: effort into material resource production and/or effort into fighting for appropriating these resources produced by others (e.g., [8–14]). This approach therefore assumes that individuals choose their behaviours freely with the goal of maximising their payoff, i.e., the rational actor model, while the material nature of the resources over which groups contest means that warfare can be regarded as a type of subsistence strategy in these formalizations. There is a parallel literature in evolutionary game theory, where instead of being freely chosen by individuals, strategies are genetically or culturally inherited (e.g., [15–18]). Both of these strands of the literature usually assume that groups are formed randomly, so that individuals mix freely between groups.

Social vertebrates and invertebrates, however, have in common that their evolution occurs in populations composed of groups of finite size with limited genetic mixing between groups [19, 20]. Such structure is relevant as it leads to interactions between genetically related individuals. Kin selection, which occurs whenever a trait expressed by an actor affects the fitness of others who are genetically related to the actor at the loci determining the trait [21–24], is therefore likely to affect the evolutionary dynamics of warfare. The literature that investigates the effects of limited genetic mixing on warfare evolution typically focuses on understanding how the demographic properties of groups, such as migration rate or sex-differences, influence the evolution of two types of behaviours: the propensity for between-groups contests and/or the fighting effort into contests [25–33]. Most of these models consider that the resources over which groups contest are reproductive in nature (e.g., mates or reproductive breeding spots). Warfare in this case should therefore be regarded as a reproductive rather than a subsistence strategy.

In this paper, we blend key elements from these different strands of the literature with the goal of contributing in two main ways. First, we decompose the individual behaviours that underpin in-group solidarity and outgroup hostility by considering the genetic co-evolution between four individual traits in a group-structured population subject to limited genetic mixing: (i) the effort into a common-pool resource within groups; (ii) the proclivity to contest other groups to appropriate their common-pool resources; and efforts into (iii) defensive and (iv) offensive structures. Distinguishing between defense and offense allows to consider potential trade-offs between the two, something that has been been somewhat neglected by previous formalizations [34]. Hence, our model can be regarded as an extension of “Guns versus butter” models, which involve a trade-off between appropriating and producing resources ([14] for review), to a situation allowing for trade-offs between appropriating, defending, and producing resources. The second way we aim to contribute is by expanding current evolutionary analyses to understand the conditions that are conducive to the emergence of polymorphism in traits underlying warfare (instead of focusing on monomorphic evolutionary equilibria like all current models). In particular, we investigate whether there are situations where peaceful individuals, characterised by low levels of belligerence and bravery (“Doves”), can coexist with hostile ones, characterised by high levels of belligerence and bravery (“Hawks”). As it turns out, our analysis suggests that such a situation can readily arise.

## 2 The Model

### 2.1 Life-cycle and evolving traits

We consider a population of asexual haploid individuals that is subdivided among *N*_g_ groups (where *N*_g_ ≫ 1 is large), each containing *N* adult individuals. All groups are subject to the same environmental conditions and are equally connected to one another by uniform random dispersal (i.e. Wright’s island model [35]). We census this population at discrete demographic time points between which the following events occur in sequence: (1) Within groups, adult individuals produce a common-pool resource that yields a material payoff (e.g. calories). (2) Each group may raid another one to appropriate its common-pool resource, leading to contests (fights) among pairs of groups over resources. (3) Each adult individual produces many offspring in quantity that depends on outcome of the previous events, i.e. on the eventual amount of resources available for reproduction after raiding (we detail this in (2.2) below). (4) Independently from one another, each offspring then disperses with a probability 0 *< m* ≤ 1 to another randomly chosen group (or remains in its natal group otherwise). (5) Finally, a randomly chosen adult dies in each group and one offspring among those competing locally is randomly chosen to fill the open reproductive spot (i.e., a Moran reproductive process [36]). Generations are thus overlapping in our model (unlike Wright-Fisher models for e.g. [36]), with individuals living on average for *N °* 1 demographic time periods. This assumption of a Moran life-cycle as well as of asexual reproduction improves mathematical tractability without loss of generality (Supplementary Material - SM for short - S5 for further discussion on this).

Against this demographic backdrop, we are interested in the co-evolution between four quantitative traits that influence interactions within- and between-groups: (1) the effort (or investment) *h* ≥ 0 made by an individual into the production of a common-pool resource for its group (helping for short); (2) the proclivity or motivation *a* ≥ 0 of an individual to raid another group, which we refer to as “belligerence”; (3) the effort *b* ≥ 0 an individual makes into group contest when its group raids another one, which we refer to as “offensive bravery”; and finally (4) the effort *d* ≥ 0 made by an individual into group contest when its group is attacked by others, which we refer to as “defensive bravery”. We assume that each of these four traits (*a, b, d* and *h*) is encoded by a separate genetic locus, at each of which we assume there is a continuum of possible alleles (i.e., the standard continuum of allele model of population genetics [37, 38]). Specifically, mutations occur during reproduction with a small probability *µ* at each locus, in which case the effect size of a mutation on the trait value is random, unbiased and weak; namely, a mutation causes a small trait deviation from the parental trait value and this deviation has mean 0 and small variance *σ*^2^.

Under these assumptions, each individual expresses a potentially unique genetically-determined trait. Thankfully, we do not need to take into account the full breadth of this variation in order to evaluate the payoff, reproduction and survival of individuals that underlay our evolutionary analysis. To that end, it is sufficient to focus here on three levels of phenotypic specification. First, we consider a *focal individual* (i.e. a representative or randomly sampled individual in the population), whose phenotype we will denote by the vector ***z***_*•*_ *=* (*a*_*•*_, *b*_*•*_, *d*_*•*_, *h*_*•*_). Second, because we will assume that individuals within groups interact in a way that can be characterised by the average trait within groups (section 2.2 for details), we do not need to specify the traits of each individual in the focal group but rather summarise this by ***z***_0_ *=* (*a*_0_, *b*_0_, *d*_0_, *h*_0_), which collects the averages of each trait among all adults of the focal group (thus including the focal individual). Finally, since mutations are rare with small effects on the phenotype, variation among individuals in the rest of the population will typically be small. We can in fact ignore the variation among- and within-groups other than in the focal group [39], and in effect consider that the rest of the population is monomorphic for the population average, which we denote by ***z*** *=* (*a, b, d, h*). In the next section, we specify under these assumptions how the evolving traits influence common-pool production within groups and raiding between groups (stages (1)-(2) stage of the life cycle in section 2.1), and in turn how this affects individual reproduction.

### 2.2 Helping, belligerence and bravery: costs and benefits

#### Common-pool production

We assume that the material payoff yielded by the common pool resource production in the focal group is given by a function *B* (*Nh*_0_), which increases in a decelerating manner with the total amount of investment, *Nh*_0_, into helping within the group (i.e. exhibiting diminishing returns; *B* (0) *=* 0, *B*′(*x*) *>* 0, and *B*″ (*x*) *<* 0 where throughout a prime ‘ denotes differentiation; and as with other relevant functions used in our model, we will later specify and explore different forms for *B* (*x*), see section 3.2 and SM S5).

#### Attacking

We assume that the probability that the focal group raids another group is given by a function 0 ≤ *α* (*a*_0_) ≤ 1, which increases with the average group belligerence (*α*(0) *=* 0, *α*′ (*a*_0_) *>* 0). Following the island model of warfare [29], we assume that when a group decides to attack another one, the group it raids is sampled randomly from the population. If two or more groups decide to raid the same group, one group is chosen at random from the attackers to perform the raid and engage into the contest for the appropriation of the common-pool resource of the raided group. As a result of these assumptions, the probability that a focal group engages into a fight over the resource of another group (i.e. decides to attack and is chosen among the attackers if there are more than one) is

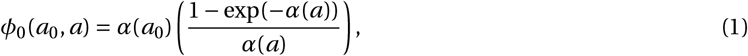

where *a* is the average level of belligerence in the rest of the population (Appendix S1 of [29] for derivation). Meanwhile, the probability that this same focal group is attacked by another group and engages locally in a contest over its own common-pool resources is

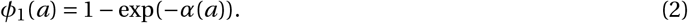

### Winning a contest

When a group raids another one, who wins the ensuing contest depends on how much the attacking group has invested into offense compared to how much the attacked group has invested into defense. More specifically, a raiding focal group with average level of offensive bravery *b*_0_ is assumed to win the contest against a raided group with average level of defensive bravery *d* with probability

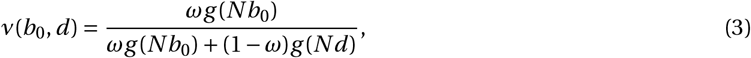

where *g* is a positive, increasing function of its argument (*g* (*y*) *>* 0, *g*′ (*y*) *>* 0). The parameter 0 *< ω <* 1 allows to tune the advantage of being offensive relative to being defensive (e.g., in the extreme case *ω =* 0, attackers always lose whereas they always win when *ω =* 1). Note that by symmetry of eq. (3), ν(*b, d*_0_) gives the probability that a focal group with average level of defensive bravery *d*_0_ loses a contest when it is attacked by another group with average level of offensive bravery *b*. In the context of conflict, eq. (3) is typically referred to as a “contest success function”, in which the choice of the function *g* allows to model different qualitative types of conflicts [8, 10, 11, 40].

### Resource distribution and fighting costs

If the attacking group wins the contest, it appropriates all the collectively produced resources of the raided group (who thus loses all its collective resources). If the attacking group loses, then both the raiding and raided groups keep their own resources. Fighting, however, is costly (for instance due to lost opportunities). We assume that the payoff available to a group is reduced by a constant −*c*_1_ *<* 0 when this group fought once (as an attacker or defender), and by −*c*_2_ *<* 0 when this group fought twice (once as an attacker and once as a defender).

### Individual payoff benefits and costs

After fighting is done, the resources that remain (if any) in a group are pooled and divided equally among its members. For example, consider the focal group with average level of helping *h*_0_ that (i) raided (and won against) another group with average investment *h* into helping, and (ii) was attacked by yet another group but also won this second fight. An individual from such a focal group will then obtain a payoff of [*B* (*Nh*_0_)*+B* (*Nh*) −*c*_2_]/*N*. Individuals also pay an individual cost due to the expression of their traits. For instance, an investment *h*_*•*_ by a focal individual to the common-pool resource will incur a cost to that individual (that increases with the actual investment *h*_*•*_). We additionally assume that both offensive (*b*_*•*_) and defensive (*d*_*•*_) bravery are costly to express, for instance due to individual resources being redirected towards offensive or defensive structures (e.g. bows, arrows or trenches). Belligerence (*a*_*•*_), by contrast, is assumed to not be associated with a direct individual cost (there are however indirect costs due to fighting captured by *c*_1_ and *c*_2_, see above paragraph). To reflect these assumptions, we write *C* (*h*_*•*_, *b*_*•*_, *d*_*•*_) for the material costs that a focal individual pays when expressing trait values *h*_*•*_, *b*_*•*_, and *d*_*•*_ (we assume these costs increase monotonically with each trait, at least linearly, i.e. *C* (0, 0, 0) *=* 0, *∂C* /*∂x >* 0 and *∂*^2^*C* /*∂x*^2^ ≥ 0 for *x* ∈ {*h*_*•*_, *b*_*•*_, *d*_*•*_}). To continue with the above example, a focal individual with traits *h*_*•*_, *b*_*•*_ and *d*_*•*_ that is a member of the focal group will then receive a payoff of [*B* (*Nh*_0_) *+B* (*Nh*) −*c*_2_]/*N* −*C* (*h*_*•*_, *b*_*•*_, *d*_*•*_).

Taking into account all possible outcomes, the expected material payoff to a focal individual can be written as

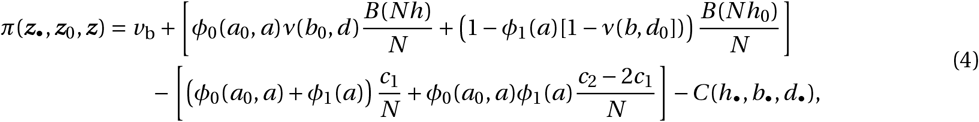

where the first term, *v*_b_ *>* 0, is some baseline payoff; the term within square brackets on the first line is the average amount of resources (over offensive and defensive contests) an individual obtains; the term within square brackets on the second line is the decrease in the amount of such resources due to the costs of contests; and the final term is the individual costs of expressing helping and bravery (see SM S1 for a derivation). When fighting probabilities are equal to one (*ϕ*_0_ *= ϕ*_1_ *=* 1) and the cost of fighting one or two fights are the same (*c*_2_ *= c*_1_ *= c*), eq. (S1) reduces to

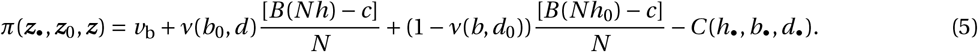

This has the same structure as the payoff function used in classical model of contests (e.g. eq. 7 of [8], eq. 1 of [14], second equation on p. 1017 of [18]), with the difference that here, a group experiences two contests: one in offense and another in defense.

### Evolutionary dynamics

We assume that the fecundity of an individual increases with its payoff. From eq. (4), it is then only a matter of bookkeeping to calculate the individual fitness of a focal individual, which depends on the payoff of other individuals in the population (SM S1 for these calculations). This lays the basis of our method of analysis of evolutionary dynamics, which is detailed in our SM S2. One can then follow our mathematical analysis from SM S3 and via an accompanying Mathematica Notebook.

## 3 Results

### 3.1 Directional selection on helping and belligerence

From our assumptions, it is clear that belligerence and bravery can only evolve by selection if other groups have produced some amount of common resource that can be appropriated by raiding. It is therefore useful to first understand the evolution of the production of the common good within groups in the absence of belligerence and bravery (*a = b = d =* 0). We find that provided helping can increase when absent in the population, then there is a unique equilibrium *h*_*§*_ for helping that satisfies the first-order condition

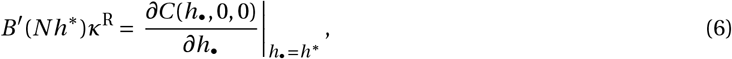

Where

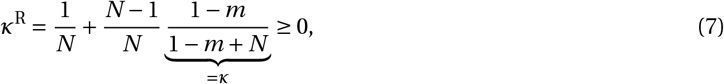

decreases with dispersal and group size (SM S3.1 for derivation). The equilibrium *h*\* defined by eq. (6) is both *convergence stable* and *locally uninvadable* under our assumptions for the benefits and cost functions (SM S2.1 for a formal definition of these terms). The population mean of helping will therefore converge to *h*^*§*^ and the phenotypic distribution will remain unimodally distributed around that mean (Suppl. Fig. 2 for e.g.). Eq. (6) says that the equilibrium investment into common pool resource production is such that the marginal cost of helping (right hand side of eq. 6) is offset by the marginal effect *B*′(*Nh*) of helping by an individual on group productivity weighted by *κ*^R^. To understand this parameter *κ*^R^, let us note first that in eq. (7), *κ* is the scaled relatedness between two individuals within groups taking kin competition into account [41, 42]. This *κ* can be interpreted as the number of units of fecundity or payoff that a focal individual is willing to forgo to increase the fecundity or payoff of a randomly sampled neighbor by one unit (for details see [43]). The parameter *κ*^R^ can then be understood as the number of such sacrificed units when the one unit increase can be obtained by any individual in the group, including the focal individual (i.e. sampled with replacement from the group, hence the superscript R). This stems from the fact that public good production benefits all individuals equally within the group, including the focal actor. Since *κ* and *κ*^R^ are both monotonically increasing functions of relatedness, they vary with demographic parameters in similar ways as relatedness (i.e. decrease with dispersal and group size, Suppl. Fig. 1). From this observation and eq. (6), we thus see that helping and common good production evolve to greater levels under directional selection when dispersal is weak and group size is small, which is a standard result across many incarnations of this problem in the literature ([41, 42] for reviews).

Let us now suppose that helping has evolved towards the equilibrium (eq. 6), but belligerence as well as bravery are absent (*a = b = d =* 0). By studying selection on belligerence in this situation (SM S3.1), we find that belligerence is favored by selection when the expected benefit from a raid exceeds its cost, i.e. when

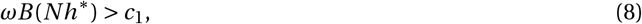

where *ω* is the probability of winning a fight when attacking when *b = d =* 0 in eq. (3). This shows that intergroup belligerence readily evolves, provided helping within groups leads to a sufficient amount of common goods that makes fighting for worth it. Once belligerence has evolved (so eq. 8 holds), this should set the stage for bravery to be favored by selection since it increases one’s chances to win a contest, either in offense or defense. We investigate this in the next section.

### 3.2 The co-evolutionary equilibrium between belligerence, bravery and helping

In order to analyse the co-evolution of all traits, we assume the following relationships: (1) 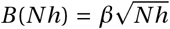 for the benefits of the common good (where *β* > 0 is a constant); (2) *α*(*a*) *= a* for the probability of attacking in a group with belligerence *a* (so that when *N =* 1, the trait 0 ≤ *a* ≤ 1 is simply the probability of attacking); (3) *g* (*b*) *= b* for the effect of bravery *b* on the winning probability; and finally, (4) *C* (*h, b, d*) *= h + c*_b_*b + c*_d_*d* for the individual cost of expressing helping and bravery, so that helping has a baseline cost of 1, and relative to this, offensive and defensive bravery have costs *c*_b_ and *c*_d_, respectively. These relationships capture the main properties of our model, while being simple enough to allow us to characterise analytically evolutionary equilibria (SM S5 for relaxation of these assumptions).

Under these assumptions, we show in SM S3.2 that there is a unique convergence stable strategy ***z***\**=* (*a*\*, *b*\*, *d*\*, *h*\*) that can be expressed as

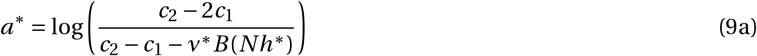

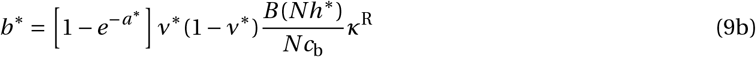

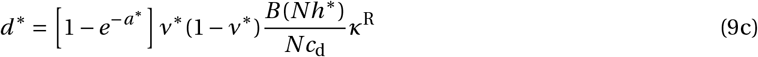

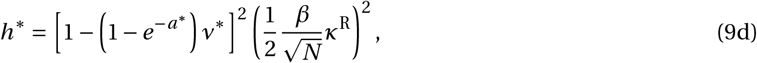

where we used

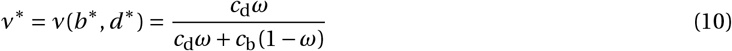

to denote the probability of winning a contest when attacking, or equivalently, of losing a contest when defending one’s own resources at the equilibrium (found by substituting the equilibria for bravery, eqs. 9b-9c, into the contest function eq. 3 with *g* (*y*) *= y*).

### Belligerence

To interpret eq. (9a), note first that *ν*\**B* (*Nh*\*) corresponds to the equilibrium expected payoff that a group receives if it raids another one (since it wins the contest with probability ν^*^ and then obtains payoff *B* (*Nh*\*)). Thus, the equilibrium for belligerence *a*\* increases with the expected group payoff of a raid and decreases with the cost of two contests relative to one contest (i.e. with *c*_2_/*c*_1_). Eq. (9a) also reveals that for *a*\* to be an interior equilibrium (i.e. 0 *< a*\**<* 1 since here *a*\* is a probability), the cost of two contests must be greater than twice the cost of a single contest (*c*_2_ *>* 2*c*_1_). If the cost of two contests is lower than this (*c*_2_ ≤ 2*c*_1_), then the cost of contests for groups that both raid and are raided is relatively low. This favours the evolution of “total” belligerence where every group attempts to raid (i.e. *a* →1).

### Bravery

To understand eqs. (9b)-(9c), we can decompose these as the product of four quantities with which both forms of bravery therefore increase: (1) the probability [1−*e*^−*a*\*^] of raiding or being raided in a population where the average belligerence is *a*\* (eqs. 1 and 2 with *a*_0_ *= a = a*\*); (2) the variance or uncertainty, *ν*\* (1‒ *ν*\*), in the outcome of a contest for a group either in offense or defense. Selection on both forms of bravery increases with this uncertainty because when one is certain to win or lose irrespective of the investment into offense or defense (e.g. when *ω =* 0 or 1), then there is no incentive for such investment; (3) the ratio of the individual benefit in the case of the group winning a contest relative to the individual cost of the relevant bravery trait, which is [*B* (*Nh*\*)/*N*]/*c*_b_ when raiding and [*B* (*Nh*\*)/*N*]/*c*_d_ when raided; and finally (4) scaled relatedness with replacement *κ*^R^ (eq. 7). We further see that when the costs of investments into offense and defense are equal (*c*_b_ *= c*_d_), individuals evolve to invest the same amount of resources into offense and defense (i.e. *b*\* *= d*\*) and that this amount is greatest when *ω =* 1/2 (so that *ν*^*^ *=* 1/2 and there is maximum uncertainty *ν*^*^ (1 ‒ *ν*\*) *=* 1/4 over outcome). The equilibrium given by eqs. (9b)-(9c) is consistent with previous models of bravery evolution for randomly mixed groups (i.e. when *m =* 1 so *κ*^R^ *=* 1/*N*, e.g. eq. 10 of [8], first equation p. 1018 of [18]), but inconsistent with those in [17, 18] concerning evolution under limited genetic mixing. The equations presented in [17, 18], however, fail a number of sanity checks (SM S3.2.2 for details).

### Helping

Finally, eq. (9d) can be understood by first recognising that the quantity *ψ* (*a*\*) *=* [1 − (1 −*e*−^*a&*\*^) *ν*\*]^2^, is the probability that a group is left with just its own common good in a population at equilibrium. This can be seen by decomposing *ψ*(*a*\*) as the product between the probabilities of two events: (1) that the group does not attack and win the ensuing contest (with probability [1 − (1 −*e*−^*a&*x002A;^) *ν*\*]); and (2) that the group does not get attacked and lose the ensuing contest (also with probability [1 − (1 − *e*−^*a&*x002A;^)*ν*\*]). In this light, eq. (9d) is intuitive. Selection for helping and participation to the common good increases with the certainty that this common good is the only source of payoff to oneself and to relatives (from *κ*^R^ in eq. 9d). Accordingly, belligerence between groups, *a*\*, reduces helping at equilibrium (as in the absence of belligerence, helping stabilises to 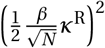, see also eq. 6).

### Payoff at equilibrium

From eq. (9), it is straightforward to obtain an explicit solution for each trait (i.e. a solution that depends only on model parameters, eq. S21 in SM S3.2.3). As expected, one characteristic feature of these solutions is that all traits at equilibrium increase with relatedness and therefore decrease with dispersal between groups (Fig. 1 A-C). What is perhaps less intuitive is that the coevolution of all four traits leads to a non-monotonic relationship between payoff at the evolutionary equilibrium and relatedness (or dispersal, Fig. 1D-F). Specifically, depending on the individual cost of offensive bravery, *c*_b_, the payoff in an equilibrium population can: (1) decrease with relatedness (for small *c*_b_, Fig. 1D); (2) first increase and then decrease in a quadratic fashion with relatedness (for intermediate *c*_b_, Fig. 1E); or (3) increase with relatedness (for high *c*_b_, Fig. 1F). This reflects the fact that payoff increases with helping within groups and decreases with belligerence between groups. As dispersal goes down and relatedness increases, selection favours more helping (which increases payoff) but simultaneously also more fighting (which decreases payoff). These antagonistic effects on payoff can lead to a situation where payoff does not monotonically increase with relatedness (in contrast to most models of social evolution). In particular, when *c*_b_ is small, belligerence tends to increase compared to helping (Fig. 1 A) causing overall a decrease in payoff (Fig. 1 D). The coevolution of belligerence and helping can therefore lead to a somewhat counter-intuitive scenario where greater relatedness and greater prosociality within groups are associated with lower payoff at equilibrium.

**Figure 1:**
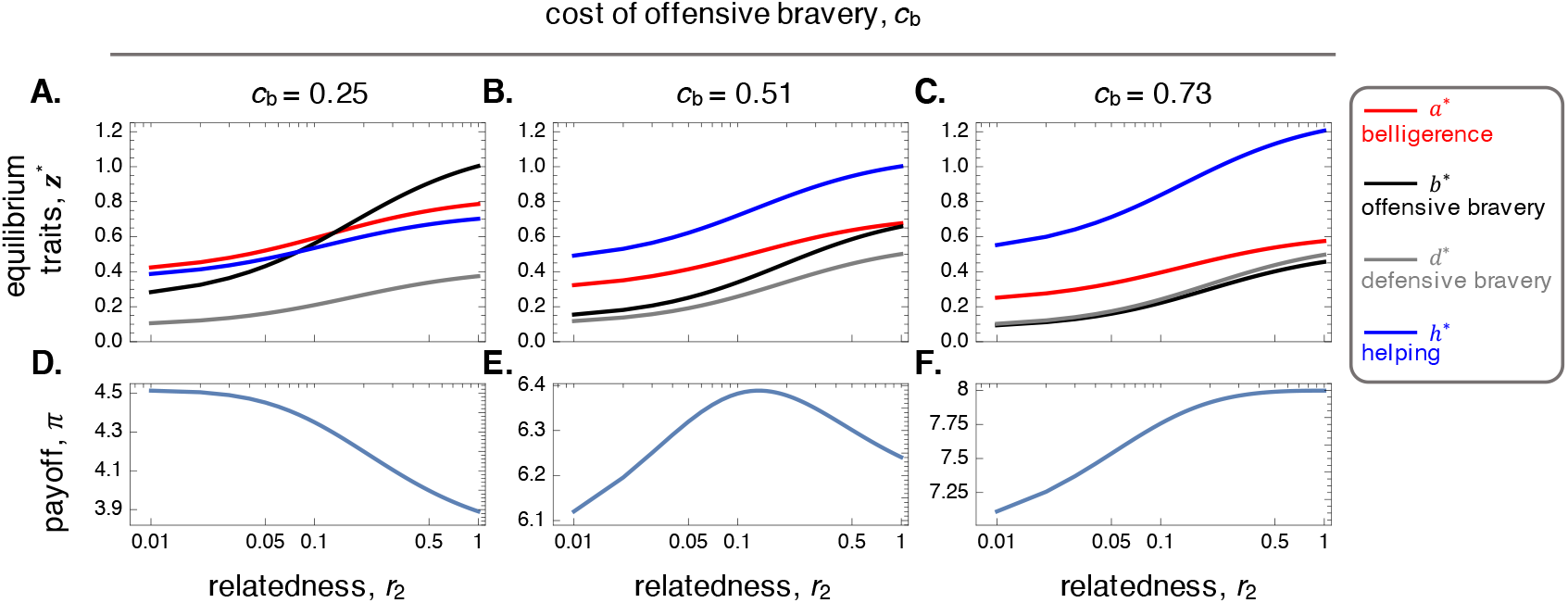
Traits *z*\* (*a*\*, *b*\*, *d*\*, *h*\*) and payoff *π*(*z*\*, *z*\*, *z*\*) at evolutionary equilibrium. **A**-**C**: Evolutionary equilibria of belligerence (*a*\* in red), offensive bravery (*b*\* in black), defensive bravery (*d*\* in gray) and helping (*h*\* in blue) as a function of pairwise relatedness (found by fixing *N =* 8 and solving *r*_2_ eq. (S12) for *m*, which is then substituted into equilibrium eqs. 9 with eqs. 7 and (S21); other parameters: *c*_1_ *=* 18, *c*_2_ *=* 115, *c*_d_ *=* 0.67, *ω =* 0.5, 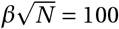, *v*_b_ *=* 0, see legend for *c*_b_). All traits increase with relatedness. **D**-**F**: Payoff as a function of pairwise relatedness (using eqs. (S1)-(S3) and traits in top row). When the cost of offensive bravery is low (*c*_b_ *=* 0.25), payoff decreases with relatedness (in D). This is because belligerence (red in A) increases more than helping (blue in A) with relatedness. By contrast, when *c*_b_ *=* 0.73, helping increases more than attack (in C) with relatedness causing a simultaneous increase in fecundity (in F).

### 3.3 The emergence and polymorphic coexistence of belligerent and pacifist individuals

#### Additive individual costs and the emergence of Hawks and Doves

Our results so far indicate that the mean phenotypes will gradually converge towards an interior evolutionary equilibrium point, provided the cost of two contests are greater than twice the cost of a single contest (*c*_2_ *>* 2*c*_1_, eq. (9)). However, we find that once such convergence has occurred, selection becomes disruptive and favors an increase in phenotypic variance, and in particular in the covariance between belligerence and offensive bravery (SM S3.2.5 for analysis). Specifically, selection favours belligerence and offensive bravery to become positively associated within individuals because genotypes that code for either more belligerence and greater investment into offense (i.e. with *a > a*\* and *b > b*\*), or less belligerence and fewer resources in offense (i.e. with *a < a*\* and *b < b*\*), have greater fitness than average when the population is at the evolutionary equilibrium. These two types gain fitness by employing two opposite strategies: to attack and win contests by investing more resources into offense, or not to attack and bypass the need to invest costly resources into offense. We respectively refer to these two types as “Hawks” and “Doves” owing to their overall phenotypic similarity to the strategies of the classical Hawk-Dove game [44, 45] as well as the nature of selection that acts upon them (see below).

We checked our mathematical analyses and investigated whether Hawks and Doves can coexist in the long run by running individual based simulations (SM S4 for procedure). Starting with a monomorphic population where each trait is absent (i.e. the value of each trait is zero), the population average of each trait rapidly converges to its predicted equilibrium (Fig. 2A). Concomitantly, the variance in belligerence also increases, with individuals gradually expressing either no or complete belligerence (*a =* 0 or 1, Fig. 2B-C). Offensive bravery *b* meanwhile also becomes polymorphic, with individuals eventually investing either some resources into offense or none at all (Fig. 2D). In contrast, defensive bravery *d* and helping *h* remain unimodally distributed so that these traits do not become significantly differentiated in the population (Fig. 2E-F). The joint equilibrium distribution of belligerence and offensive bravery (Fig. 3A) confirms our analytical predictions that these two traits become positively associated, and further reveals that highly-differentiated Hawks and Doves coexist in the long run (respectively the top right and bottom left clusters in Fig. 3A). By contrast, there is no clearly discernible association among any other pair of traits, so that Hawks and Doves both express helping and defense bravery in about the same amount (Fig. 3B-F). These weak associations are confirmed by the weak equilibrium covariances among traits other than between belligerence and offensive bravery (Fig. 3G).

**Figure 2:**
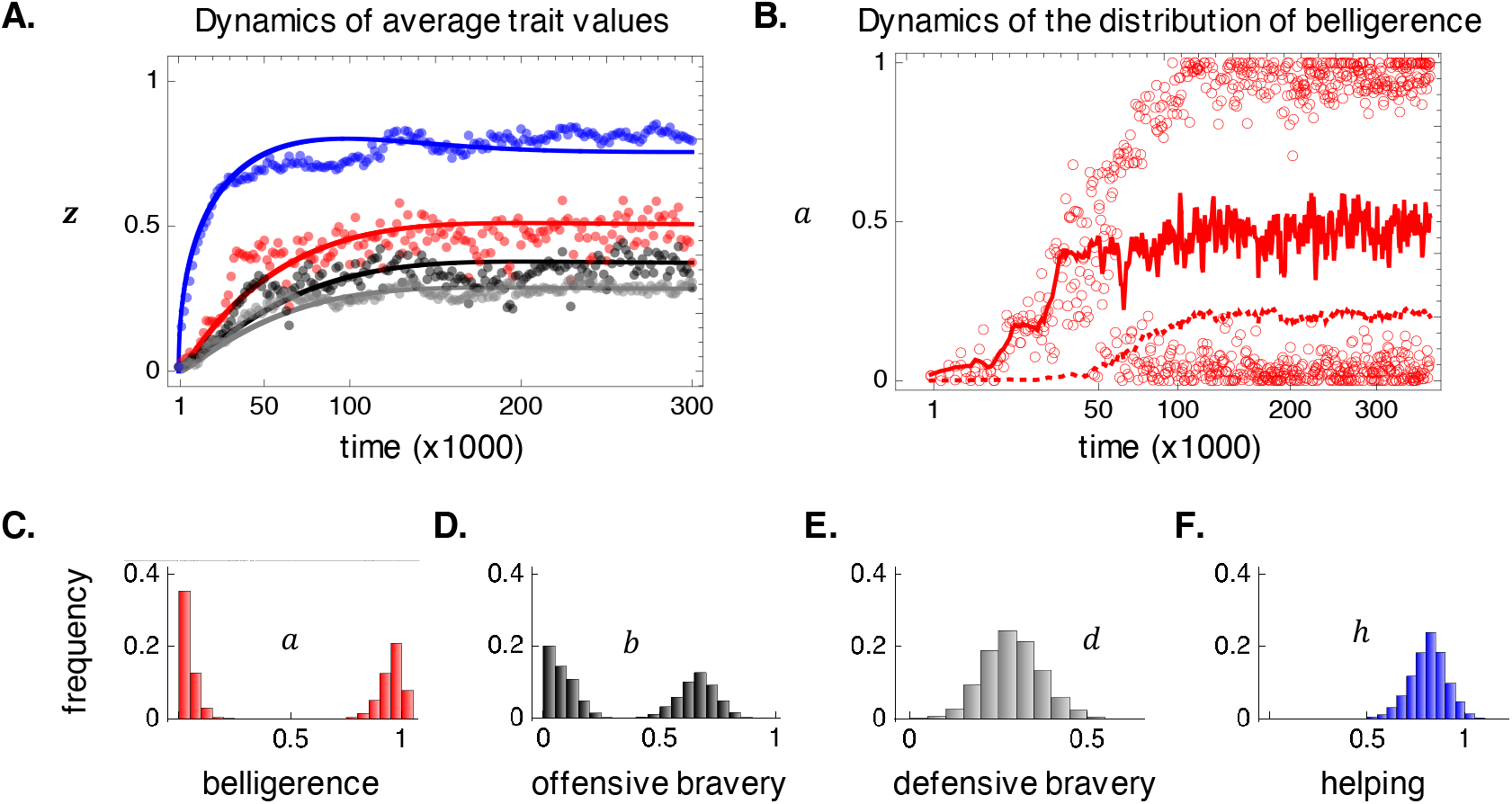
The emergence of polymorphism. **A**: Average trait values in the population as a function of time (with a unit of time corresponding to an iteration of the life cycle) in a population where all traits are initially absent (*a*_*t*_ in red; *b*_*t*_ in black; *d*_*t*_ in gray; *h*_*t*_ in blue; observed in a simulation in dots, SM S4 for simulation details; analytically predicted in full lines, from eq. (S9) with variance-covariance matrix **G** *= δ* ^2^**I** where **I** is the identity matrix and *δ =* 0.043 was chosen heuristically; *N =* 8 and *m =* 0.467 so that *r*_2_ *=* 0.125; otherwise same parameters as Fig. 1 middle). **B**: Individual values of belligerence observed in a simulation (empty circles, shown for two individuals randomly sampled every 800 time points, same parameters as A), as well as the observed trait population average (full line) and variance (dotted line, time on a log scale). Note that polymorphism occurs before the population average has completely converged to its equilibrium in simulations due to mutations being relatively large (to speed up computation time). But as predicted, the trait variance starts to increase significantly only once the population average has stabilised. **C**-**F**: Distribution of each trait in a simulation (calculated from time 250’000 for 100’000 time steps, same parameters as A).

**Figure 3:**
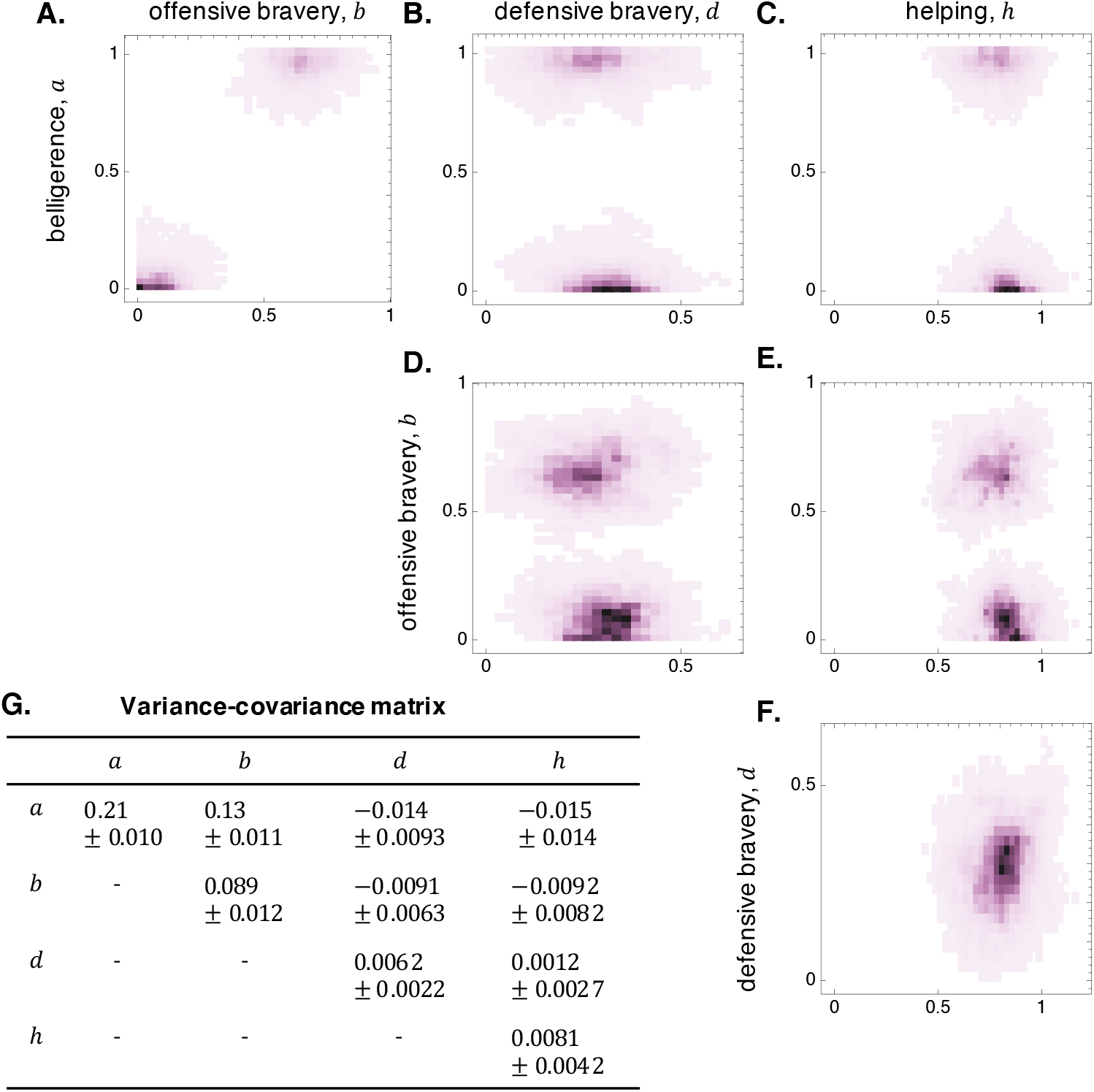
Correlations among traits. **A**.-**F**. Joint distribution of each pair of traits in a simulated population at equilibrium (calculated from time 250’000 for 100’000 time steps, same simulation as Fig. 2, darker shade means greater density). **G**. Mean *±* standard deviation of the variance-covariance matrix evaluated in the same simulated population and time points as A-F; variances are on the diagonal while covariances among each pair traits are on the off-diagonal (see SM eq. (S8) for definition).

These analyses show that the evolution in our model leads to the gradual emergence and maintenance of highly differentiated-types owing to negative frequency-dependent interactions. The pattern of frequency-dependence can be understood by focusing on Hawks. These are particularly successful when they are rare as they tend to engage their group into a raid and to win contests against groups consisting mainly of Doves. But as Hawks become common, their groups suffer from attacks by other groups consisting mainly of Hawks, and as a result pay the high cost of two contests rather than one (i.e. pay *c*_2_ rather than *c*_1_, where *c*_2_ *>* 2*c*_1_). Passed a critical frequency, Hawks thus become less successful than Doves, allowing both types to be maintained in the population. This frequency-dependence is thus reminiscent to that of the Hawk-Dove game [44, 45], with interactions here mediated by group-structure rather than occurring strictly among individuals.

#### Polymorphism within and between groups

The group structure of the population raises the question whether coexistence occurs within or between groups, in other words whether groups tend to be composed of one type (only Hawks or only Doves) or a mix of both. The distribution of belligerence in a single focal group over time suggests the latter (Fig. 4A), with the group very often consisting of both types. Nevertheless, when we compare the average level of belligerence within a focal group to the population average (dots vs full lines in Fig. 4B), we see that the focal group experiences significant variation over time. In other words, even though both Hawks and Doves co-occur within a group, one morph will typically dominate at any given time. Groups will therefore tend to be differentiated according to whether they are composed of more or less of one type. We can quantify this at the level of the population by calculating the phenotypic differentiation among groups at the belligerence locus (Fig. 4C, red). We find that this differentiation is no different before and after the polymorphism emerged or to differentiation at the helping locus (which recall never becomes clearly polymorphic, Fig. 4C, blue). In fact, differentiation among groups at the belligerence locus is the same as expected for a neutral locus (Fig. 4C, black). This shows that negative frequency-dependent selection does not generate significant departures in morph distribution among groups compared to neutral expectation. Put differently, how the two different morphs are distributed among groups is primarily determined by dispersal and local genetic fluctuations (due to variance in reproductive success that generates identity-by-descent) and thus reflect the pattern of relatedness.

**Figure 4:**
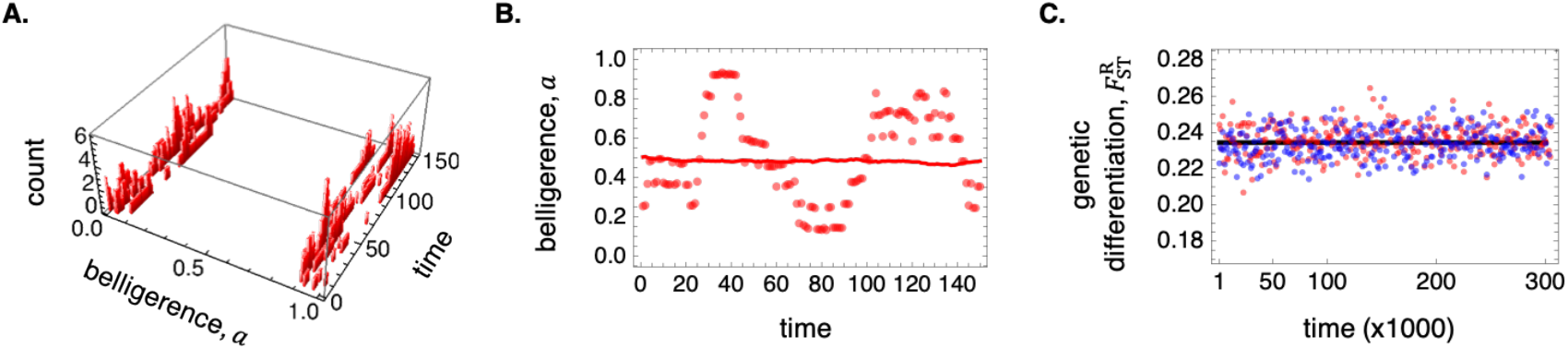
Polymorphism within and between groups. **A**. Distribution of belligerence (number of individuals) in a focal group over time (150 steps, same simulation as in figure 2 from time 299’000 onwards). **B**. Focal group-(dots) and population-(full line) average belligerence (same group as in A). **C**. Genetic differentiation among groups 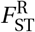 (*F*_ST_ with replacement, i.e. the ratio of the variance among groups of group-averages to the total trait variance in the population) over time at the belligerence (in red) and helping (in blue) loci in a simulated population, against neutral expectation (in black full line, calculated from 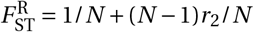 where *r*_2_ is the neutral relatedness coefficient, eq. S12, same simulation as in figure 2).

We additionally examined the effects of different levels of dispersal on polymorphism (i.e. different values of *m*). Although simulations generally show significant phenotypic variation at the belligerence and offensive bravery loci (Suppl. Fig. 3A-C), differentiation among the Hawk and Dove morph is clearest and most stable where relatedness within groups is high (Suppl. Fig. 3D). This reflects that much of the frequency-dependent selection in our model is due to interactions between groups. As a result, selection is most able to discriminate among morphs and therefore favour their differentiation when morphs are unequally distributed among groups, which as we saw in the preceding paragraph happens when relatedness is non-zero. In fact, polymorphism among Hawks and Doves is weak if non-existent under complete random group mixing (and relatedness is zero, Suppl. Fig. 3A & D).

#### Non-additive costs and the emergence of Scrounging Hawks and Producing Doves

One assumption we have made so far is that the individual costs of investing personal resources into bravery (offensive and defensive) and helping are additive (within and between traits, i.e. that *C* (*h*_*•*_, *b*_*•*_, *d*_*•*_) *= h*_*•*_ *+ c*_b_*b*_*•*_ *+ c*_d_*d*_*•*_). Complementarity or antagonistic effects between traits (i.e. when different traits respectively have positive or negative non-additive effects) on individual costs can significantly influence how selection shapes associations between social traits and therefore on the nature of adaptive polymorphism when fitness depends on multiple traits [e.g. 39, 46–48]. In the context of inter-group contests and intra-group helping, one relevant scenario to investigate is where bravery traits complement one another, for instance because weapons or behaviours that are useful in offense are also useful in defense, but antagonistic with helping, for e.g. because characteristics that are beneficial in situations of conflict are counter-productive in prosocial situations. One way to capture this scenario is to implement non-additive individual costs in eq. (4) of the form

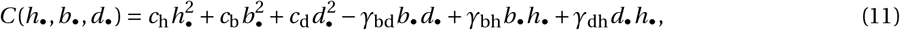

where *γ*_bd_ *>* 0 controls the complementarity among offensive and defensive bravery (so that individual costs are lower when one unit of resource is invested in both types of bravery compared to two units invested in a single type), while *γ*_bh_ *>* 0 and *γ*_dh_ *>* 0 tune the antagonism between each respective bravery trait and helping.

Numerical exploration of the mathematical model using eq. (11) suggests that polymorphism still emerges in this case and that it is still driven by the coevolution between belligerence and offensive bravery (Mathematica Notebook). However, phenotypic associations now involve more traits. In particular, selection now favours a negative association between helping with offensive and defensive bravery. Individual-based stochastic simulations confirm this and further highlight the maintenance of a negative association among belligerence and helping (Fig. 5). Specifically, the Hawk morph is now also characterised by little within-group helping (morph on the top left of Fig. 5A), and the Dove morph by high levels of helping (morph on the bottom right of Fig. 5A). In fact, all other traits are now negatively associated with helping (last column of Fig. 5B). This is due to the extra individual costs suffered by individuals that combine helping with any form of bravery. Since Hawks are characterised by high levels of bravery, they evolve lower helping because of these extra costs. Doves, meanwhile, can continue to invest resources in helping since they invest little or no resources into bravery.

**Figure 5:**
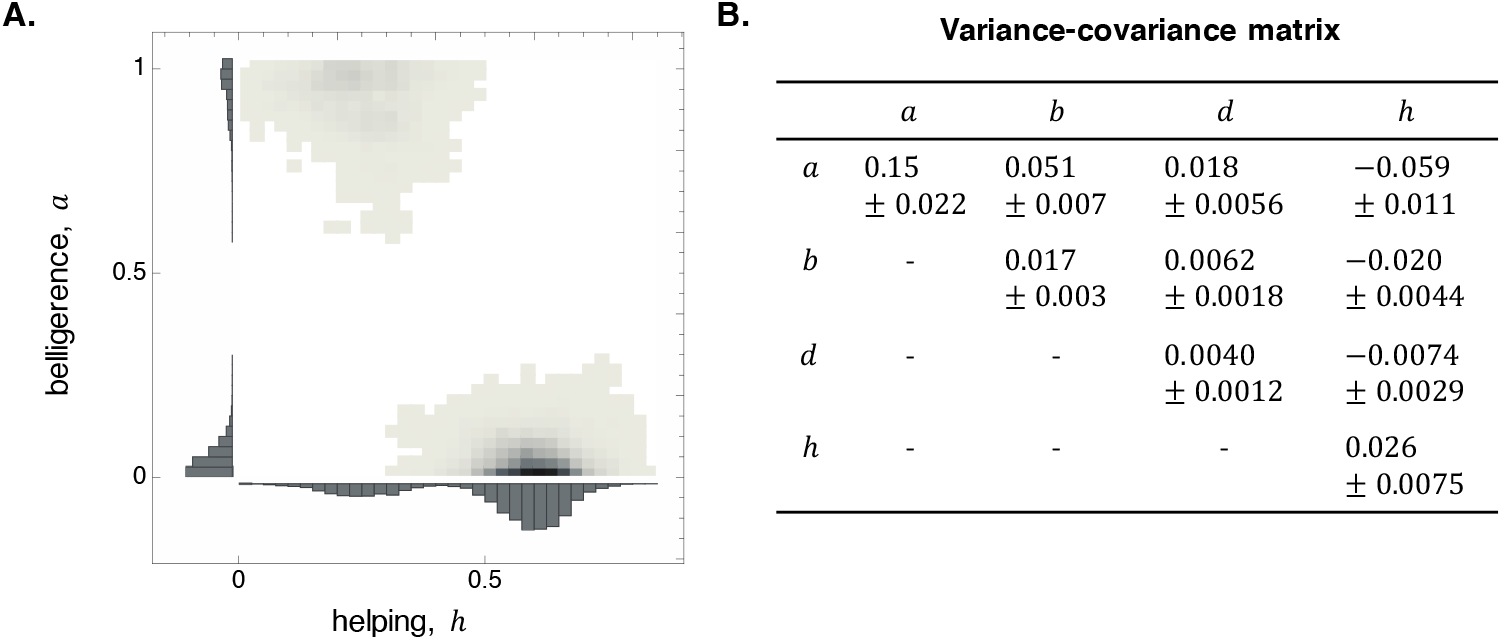
Negative association between helping and contest traits. **A**. Joint distribution of belligerence and helping in a simulated population at equilibrium, where offensive and defensive bravery have complementarity effects among each other but antagonistic effect with helping (at the individual level), i.e. where *C* (*h*_*•*_, *b*_*•*_, *d*_*•*_) is given by eq. (11) (with *c*_b_ *=* 0.7, *c*_d_ *=* 1.3,*c*_h_ *=* 1, *γ*_bd_ *=* 0.5, *γ*_bh_ *=* 2.2, *γ*_dh_ *=* 2.2; other parameters: same as Fig. 2; joint distribution calculated over 100’000 time points after 150’000 of evolution). **B**. Mean *±* standard deviation of the variance-covariance matrix evaluated for the same simulated population as A. In contrast to Fig. 3G, the covariance among helping (*h*) and belligerence (*a*) is now significantly different to zero and negative. And while this covariance may seem small, the correlation among belligerence (*a*) and helping (*h*) is large (as shown in **A**).

These results show how complementarity and antagonistic effects among traits can lead the polymorphism in our model to become more complex and involve further traits (such as helping). The main characteristics of the Hawk and Dove morphs, however, have not changed due to such non-linear effects, with some individuals with a strong preference for raiding and investing resources into offense, and others favouring not to raid and investing no resources into offense. We test and discuss the robustness of this polymorphism further in our SM S5, where in particular we investigate the effects of various contest functions (via *g* (*y*)), group decision making (via *α*(*a*)) and benefit functions (via *B* (*y*)). We find that in all examined cases, Hawks and Doves are still expected to emerge as in our baseline model.

## 4 Discussion

Our results indicate that selection can readily lead to the emergence of a social polymorphism where two highly-differentiated morphs relevant to warfare eventually co-exist: one that codes for belligerence and offensive bravery (Hawks) and the other for neither (Doves, Fig. 3). The frequency-dependent interactions that maintain this polymorphism are close to those characterising the classical Hawk-Dove game [44, 45], where Hawks are favored when rare but disfavored when common as they engage with other Hawks and suffer an extra (non-additive) cost due to fighting. This extra cost is typically captured in the classical Hawk-Dove game by the condition that the cost *C* for a Hawk to lose against another Hawk is greater than the value *V* of the resource obtained in case of a win (i.e. *C > V* ; whereas in the additive case, *C = V* which would disfavor Doves, always). Similarly, one prerequisite for frequency-dependent interactions to lead to co-existence in our model is that the cost of contests is greater than additive (with the cost of two contests *c*_2_ greater than twice the cost of a single contest *c*_1_, *c*_2_ *>* 2*c*_1_, eq. S21 in SM S3). These extra costs prevent groups of Hawks to dominate at all frequencies and allow Doves to thrive when rare.

There are nonetheless also significant differences between the classical Hawk-Dove game for inter-individual conflicts and our inter-group conflict interaction scenario. First, the two morphs that are held in stable polymorphism in our model are not preexisting, but rather emerge from disruptive selection and gradual evolution. In the same way that polymorphism does not emerge when belligerence evolves alone in our model (SM S3.2.5), the mixed strategy characterized by the probability of taking the action “Hawk”, usually understood as “Attack”, also converges towards an equilibrium point and does not experience disruptive selection in the classical Hawk-Dove game [49]. This brings us to a second difference: the strategies that the two coexisting types employ at equilibrium in our model are more complex than simply playing “Attack” or not. Rather those strategies consist of composite behaviours including the propensity to raid and investment into offensive abilities. Interestingly, a similar polymorphism has been found to emerge in well-mixed populations with interindividual conflicts (as in the Hawk-Dove game), and where the probability of playing Hawk coevolves with a physiological trait that is costly to express but that increases the probability of a win against another Hawk [50]. Like in our model, it is the coevolution of these two traits that leads to polymorphism emergence. One further conceptual difference regarding strategies is that Hawks always beat Doves in the classical game, whereas here, individuals of low belligerence nevertheless also invest significant resources into defense. Accordingly, groups consisting mainly of Doves in our model have a non-zero probability of winning against raiding groups consisting of mostly Hawks.

But perhaps the most significant way that we depart from the classical Hawk-Dove (as well as [50]), is that frequency-dependent interactions are mediated through group-structure in our model. Rather than lone individuals, it is groups with a majority of Hawks that tend to partake in raids and groups with a majority of Doves that tend not to. So even though most groups are composed of a mix of both morphs due to dispersal (Fig. 4A), the maintenance of polymorphism through negative frequency-dependent interactions relies on variation in this mix among groups (Fig. 4B-C for e.g.). Accordingly, when groups are formed via complete random mixing (i.e. complete dispersal, *m =* 1 and relatedness is zero) or group size is infinitely large (*N* → ∞), selection is unable to discriminate among morphs and cause their differentiation since groups show very little variation in their composition (Suppl. Fig. 3A). By contrast, where relatedness is positive, highly-divergent morphs can be observed (Suppl. Fig. 3B-D). In fact, limited dispersal tends to stabilise the polymorphism (Suppl. Fig. 3D), with selection remaining disruptive even where dispersal is severely limited (i.e. very close to *m =* 0, Mathematica Notebook). This differs also from previous models where frequency-dependent interactions happen only among individuals of the same group, in which limited dispersal and thus relatedness inhibits disruptive selection [as it reduces the amount of local genetic variation and thus differentiation within groups, 39, 51–53].

In spite of these group-effects, one should keep in mind that selection occurs at the level of the gene (or replicator) and that these are expressed by individuals. As a result, where the individual fitness costs of the different traits interact with one another, evolutionary dynamics can lead to more traits becoming associated to the social polymorphism. In particular, if helping is antagonistic with offensive and defensive bravery while both forms of bravery are complementary (eq. 11), then the two co-existing morphs we observe consist of individuals that either do not participate in common-pool resource production but only in its defense and appropriation (“Scrounger Hawks”); or only invest into common pool resource production (“Producer Doves”, Fig. 5). The negative frequency-dependent interactions that maintain these two morphs are then reminiscent to those characterising the classical Scrounger-producer game [54]. Beyond this scenario, our results suggest that through fitness interactions with bravery traits, other relevant social traits (such as the tendency to lead or follow) may become associated to the social polymorphism we have described.

The apparent ease with which polymorphism emerges in our model raises the question why it has not been reported in previous papers that have investigated the (genetic) co-evolution of traits involved in warfare [26, 28– 31, 33]. By comparing these models to ours, we find that this is likely due to divergent formulations for fitness (compare eqs. S5 and S6 in SM). This divergence comes from our perspective of warfare as a subsistence strategy, whereas previous papers allowing for the coevolution of belligerence and bravery considered it as a reproductive strategy (but see [30]). These models typically assume that (i) groups loosing contests are repopulated (partially or completely, or their females mated) by winning groups, and (ii) that belligerence has fixed or unconditional costs to the individual that expresses it (in contrast to our model where belligerence has conditional costs, which are incurred only if raids take place). The direct extra costs associated with greater belligerence under such assumptions make it more difficult for Hawks to differentiate from the population in these models compared to ours and therefore for polymorphism to emerge (SM S1.2.2 and S3.2.5 for a more formal explanation). But nothing in principle precludes from considering conditional costs in models of war-fare as a reproductive strategy and thus allow for polymorphism. Meanwhile, models that consider warfare as a subsistence strategy typically ignore belligerence and focus on bravery evolution (e.g., [15–18]). But as we have established, this coevolution between belligerence and bravery is necessary to the emergence of polymorphism.

To sum up, we have proposed a model of warfare evolution in which an initially asocial and undifferentiated population evolves towards within-group solidarity and between-group hostility enacted by highly-differentiated individuals. This as long as group size is not too large and dispersal is limited. Indeed, with complete dispersal, no differentiation occurs in our model, and with large group size, no social trait evolves to begin with as the selection pressure on each trait is of the order of the inverse of group size (1/*N*). This effect of group size on the strength of selection is, provided a few exceptions, common to all models of evolution of prosocial traits affecting group members indiscriminately [55], and thus applies to those aforementioned on warfare evolution as recently illustrated in simulations [33]. These models and ours are therefore most relevant to small-scale societies (or populations with small local *effective* size and *effective* migration rate, e.g., eq. 9. 59 in [24]). For such societies, our model suggests that between-group aggression can be a potent mechanism for the maintenance of within-group trait diversity and behavioral syndromes, in particular favouring a positive association between belligerence and bravery.

## Supporting information

Supplementary material S1-S5, figures, matehmatica notebook and code.

## Acknowledgements

We thank Carsten De Dreu and Zegni Triki for their invitation to contribute to this special issue and for their comments on a previous version of this paper, as well as three anonymous reviewers.

