## Supplementary material S1-S5, figures, matehmatica notebook and code. for "Evolution of warfare by resource raiding favors polymorphism in belligerence and bravery": BioRxiv_SuppMat.pdf

### Supplementary Material for "Evolution of warfare by resource raiding favors polymorphism in belligerence and bravery"

#### S1 Payoff and fitness

In this first supplement, we derive the expected payoff that a focal individual obtains (in S1.1) from which we then characterise its fitness (in S1.2).

##### S1.1 Expected payoff

From the assumptions and notations of the main text (section 2.2), it follows that the expected material payoff to a focal individual with traits $\boldsymbol{z}_\bullet$ in a group with average trait values $\boldsymbol{z}_0$ when the population average trait values are $\boldsymbol{z}$ can be written as

$$\pi(\boldsymbol{z}_\bullet,\boldsymbol{z}_0,\boldsymbol{z}) = v_\text{b} + \phi_0(a_0,a)\pi_\text{a}(\boldsymbol{z}_0,\boldsymbol{z}) + (1-\phi_0(a_0,a))\pi_\text{b}(\boldsymbol{z}_0,\boldsymbol{z}) - C(h_\bullet,b_\bullet,d_\bullet), \tag{S1}$$

where $v_\text{b} > 0$ is some baseline payoff, $\pi_\text{a}(\boldsymbol{z}_0,\boldsymbol{z})$ is the expected payoff to the focal individual, conditional on its group engaging into a raid [which occurs with probability $\phi_0(a_0,a)$], and $\pi_\text{b}(\boldsymbol{z}_0,\boldsymbol{z})$ is the expected payoff to the focal, conditional on the focal group not raiding another group [which occurs with probability $1-\phi_0(a_0,a)$]. Let us first consider $\pi_\text{b}(\boldsymbol{z}_0,\boldsymbol{z})$ as it is simpler to obtain. Given that the focal group has not raided another one, the expected payoff to the focal individual is

$$\pi_\text{b}(\boldsymbol{z}_0,\boldsymbol{z}) = \phi_1(a)\left[-\frac{c_1}{N} + (1-v(b,d_0))\frac{B(Nh_0)}{N}\right] + (1-\phi_1(a))\frac{B(Nh_0)}{N}, \tag{S2}$$

which can be understood as follows. With probability $\phi_1(a)$, the focal group is attacked. In this case, the focal individual necessarily pays a cost $-c_1/N$ for one fight but only gets to retain its share $B(Nh_0)/N$ of the common resource if its group wins the fight against its raiders, which occurs with probability $1-v(b,d_0)$. With probability $(1-\phi_1(a))$, the focal group is not attacked and therefore the focal individual always gets its share $B(Nh_0)/N$. Using similar arguments, we find that the expected payoff of the focal individual given that its group has participated in a raid is

$$\pi_{\mathrm{a}}(z_0,z)=v(b_0,d)\left\{\frac{B(Nh)}{N}+\phi_1(a)\left[-\frac{c_2}{N}+(1-v(b,d_0))\frac{B(Nh_0)}{N}\right]+(1-\phi_1(a))\left[\frac{-c_1+B(Nh_0)}{N}\right]\right\}$$
$$+(1-v(b_0,d))\left\{\phi_1(a)\left[-\frac{c_2}{N}+(1-v(b,d_0))\frac{B(Nh_0)}{N}\right]+(1-\phi_1(a))\left[\frac{-c_1+B(Nh_0)}{N}\right]\right\}, \quad \text{(S3)}$$

where the first line consists of the probability $v(b_0,d)$ that the focal group wins the raid it engaged in, multiplied to the expected payoff in this case (between curly brackets). Conversely, the second line is the probability $1-v(b_0,d)$ that the focal group loses the raid it engaged in multiplied by the relevant payoff obtained in that case (between curly brackets also). Substituting eqs. (S2)–(S3) into eq. (S1) and re-arrangements yield eq. (4).

#### S1.2 Fitness

We assume that an individual's fecundity, i.e. the number of offspring produced during stage (3) of the life cycle, increases with its expected payoff according to a function $F$, so that the fecundity of a focal individual with payoff $\pi$ is written as $F(\pi)$. We assume that this function is positive and decelerating with expected payoff (i.e. $F'' \leq 0$). Formally, the fecundity of an individual should really be a function of its *realised* payoffs, which depend on a specific sequence of events (such as whether fighting took place, whether the fight was won and so on), rather than *expected* payoffs, which are averaged over all possible outcomes (as in eqs. S1-S3). In writing fecundity directly as a function of expected payoffs only, we are essentially assuming that the deviation between realised and expected payoffs is small (specifically, with $\Pi$ denoting the random variable for the payoff to the focal individual, we ignore terms of order $\mathrm{E}[(\Pi-\pi)^2]$ and higher, where $\mathrm{E}[\cdot]$ stands for the expectation operator over all relevant stochastic effects). This is a standard albeit often left unspecified assumption in evolutionary game theory. One alternative to such an assumption is to define payoff directly in units in fecundity, in which case payoff is simply fecundity (i.e. $\pi = F(\pi)$). In any event, once we have specified its fecundity we can determine the *fitness* of an individual, which is its expected number of surviving offspring produced over one full iteration of the life-cycle (e.g., [1, 2]) and lays the foundation of our evolutionary analysis. We detail such fitness function for our model, which considers warfare as a subsistence strategy, below (in S1.2.1), as well as the typical fitness function that characterises models where warfare should rather be considered as a reproductive strategy for contrast (in S1.2.2).

##### S1.2.1 Warfare as a subsistence strategy

In order to obtain the expression for the fitness of a focal individual under the assumptions of our model (section 2.1), let us first label the other individuals in the focal group (i.e., the focal's neighbours) as individual "2", "3" until "$N$", and denote their respective traits as $z_2$, $z_3$ and so on, whereby we can write the average trait in the focal group as

$$z_0=\frac{z_\bullet+z_2+z_3+\sum_{j=4}^{N}z_j}{N}. \quad \text{(S4)}$$

It will also be useful to collect all the trait values of the neighbours of the focal into the vector $z_{-\bullet}=(z_2,z_3,\ldots)$. Then, according to the life-cycle detailed in section 2.1, the fitness of the focal individual is

$$w(z_\bullet,z_{-\bullet},z)=\frac{N-1}{N}+\frac{(1-m)F(\pi(z_\bullet,z_0,z))}{(1-m)[F(\pi(z_\bullet,z_0,z))+\sum_{j=2}^{N}F(\pi(z_j,z_0,z))]+mNF(\pi(z,z,z))}+\frac{mF(\pi(z_\bullet,z_0,z))}{NF(\pi(z,z,z))}, \quad \text{(S5)}$$

where recall $\pi$ is the payoff function (eq. S1) and $F(\pi)$ is the fecundity of an individual with payoff $\pi$. Equation (S5) can be read as the sum of three fitness components. (1) The first summand is the probability that the focal survives. (2) The second summand is the probability that the open philopatric spot is filled by one of its offspring. This consists of the ratio of the number of the focal's offspring that remain philopatric to the total number of offspring that enter competition in the focal patch (comprised of all those that remain philopatric – factored by $(1-m)$ – and all those that disperse from other patches – factored by $m$). (3) The last summand is the expected number of spots filled in other patches, given by the ratio of the number of the focal's offspring that disperse to the expected total number of offspring that compete in another patch.

Equation (S5) has the standard form of individual fitness under the island model of dispersal coupled with a Moran process (Box 1 in [3]). It could be straightforwardly expanded to consider the case where payoffs affect survival rather than fecundity [3]. Note also that to obtain the expression for fitness under a Wright-Fisher process (i.e. where all individuals are replaced per life-cycle iteration), one simply removes the first summand of eq. (S5) and multiplies the rest by $N$. More generally, assuming that the spoils of warfare only influence fecundity or survival, i.e. where warfare is conceived as a subsistence strategy, eq. (S5) can easily be amended to consider other common variations, such as where regulation occurs before dispersal, or where individuals survive with fixed probability.

###### S1.2.2 Warfare as a reproductive strategy

In order to highlight the key difference in fitness between co-evolutionary models where warfare is a subsistence (as in eq. S5) vs. a reproductive strategy (as in [4–9]), let us consider a simple representative case of the latter under the following assumptions: that the winning group takes over all the breeding spots of the losing group; that density-dependent regulation of offspring occurs before dispersal (i.e. soft selection); and that there is no difference between offensive and defensive bravery ($b = d$). In this case, the fitness of a focal individual can be written as

$$w(\boldsymbol{z}_\bullet, \boldsymbol{z}_0, \boldsymbol{z}) = \left[1 + \underbrace{\phi_0(a_0, a)v(b_0, b)}_{\text{gaining a patch}} - \underbrace{\phi_1(a)v(b, b_0)}_{\text{loosing own patch}}\right] \times \frac{1 - C(a_\bullet, b_\bullet)}{1 - C(a_0, b_0)} \tag{S6}$$

(obtained from eq. 11 of [6] by setting $m = 0$ and $h = 0$). The main differences between eq. (S5) and eq. (S6) is that for the latter: (i) there is no common pool resource production; (ii) the cost of belligerence is fixed, i.e., it now appears along the other traits in the cost function $C(a_\bullet, b_\bullet)$ (in contrast to eq. S1 where belligerence has conditional costs, $c_1$ and $c_2$); (iii) the benefits of warfare and the costs of trait expression are multiplicative (i.e., the benefits of warfare, which are in the square brackets, multiply the costs, while in eq. S1 the benefits of warfare and these costs of trait expression add up); and finally, (iv) the gains of warfare will not be partly destroyed by competition, i.e., the gains of warfare affect the denominator in eq. (S5) but not in eq. (S6). Having these differences in mind is useful to contrast our results with those of previous studies (see section S3.2.5).

#### S2 Mathematical evolutionary approach

In this supplement, we outline our mathematical analysis for the joint evolution of the four traits of interest ($a$, $b$, $d$ and $h$). It is based on an evolutionary quantitative genetics and adaptive dynamics model for group-structured populations that is tightly connected to invasion analysis [10]. This model tracks the dynamics of a multi-trait phenotypic distribution, assuming that the processes of selection and mutation are such that this distribution is approximately multi-variate Gaussian across the whole population (i.e. over all individuals and all groups) with small (co)variance (specifically, assuming that the largest absolute value among all traits' (co)variances can be written as $\delta^2$ where $0 < \delta \ll 1$ is a small parameter). Note that this assumption does not require that the realized distribution of phenotypes within a focal group at any given demographic time period is Gaussian, but rather that its time average is. Such assumption of normality has been shown to give accurate predictions for the evolution of traits' means and (co)variances, even where selection generates significant deviations from normality (refs. [11] for well-mixed, and [10] for dispersal-limited populations).

With the assumption of multivariate normality, the phenotypic distribution at any time $t$ for our warfare model can be described by the vector of means,

$$\boldsymbol{z}_t = \begin{pmatrix} a_t \\ b_t \\ d_t \\ h_t \end{pmatrix}, \tag{S7}$$

and the (symmetric) variance/covariance matrix

$$\mathbf{G}_t = \begin{pmatrix} G_{aa,t} & G_{ab,t} & G_{ad,t} & G_{ah,t} \\ G_{ba,t} & G_{bb,t} & G_{bd,t} & G_{bh,t} \\ G_{da,t} & G_{db,t} & G_{dd,t} & G_{dh,t} \\ G_{ha,t} & G_{hb,t} & G_{hd,t} & G_{hh,t}, \end{pmatrix} \tag{S8}$$

where $G_{uv,t} \sim \mathcal{O}\left(\delta^2\right)$ is the genetic covariance among traits $u$ and $v$ at time $t$ (so that $G_{uv,t} = G_{vu,t}$). Hence the dynamics of the phenotypic distribution are given by the joint dynamics of the vector $\boldsymbol{z}_t$ and the matrix $\mathbf{G}_t$. However, as we detail below, we do not need to consider these dynamics jointly when $\delta$ is small.

#### S2.1 Evolution in two time scales

In brief, the upshot of the analysis we follow is that when mutations are rare and have small effects on phenotypes (so that $\delta$ is small), the evolutionary dynamic can be decomposed into two time scales. First, the population evolves under directional selection whereby the average trait values $\boldsymbol{z}_t$ in the population change gradually, but the variance in each trait and covariance among each pair of traits remains small and approximately constant (i.e., $\boldsymbol{z}_t$ changes while $\mathbf{G}_t$ can be held constant). Once the population average has converged to an equilibrium for directional selection, a so-called "convergence stable strategy" (that is thus an attractor of the evolutionary dynamics), selection shapes the traits' (co)variances (i.e. $\mathbf{G}_t$ changes while $\boldsymbol{z}_t$ remains fixed for its equilibrium). An analysis of selection close to convergence stable strategies then allows to establish whether selection is (a) stabilising, keeping traits' (co)variances small so that the phenotypic distribution remains unimodal and centred around the equilibrium which is thus "locally evolutionary stable" or "uninvadable" (so that $\mathbf{G}_t$ converges); or (b) disruptive, favoring an increase in the variance (and possibly covariance) of some traits (so that $\mathbf{G}_t$ diverges). Due to disruptive selection, the population may eventually undergo "evolutionary branching" [12], whereby the phenotypic distribution becomes multi-modal so that two or more clearly differentiated morphs emerge. These morphs may differ in multiple traits owing to correlational selection, which favours specific associations between traits within individuals. We detail mathematically the two time scales of evolutionary dynamics and corresponding effects of selection in the next two sections (directional selection in S2.2 and stabilising/disruptive selection in S2.3).

#### S2.2 Directional selection

**The selection gradient.** First, the population evolves under directional selection. To the leading order in $\delta$, the change $\Delta \boldsymbol{z}_t = \boldsymbol{z}_{t+1} - \boldsymbol{z}_t$ in average trait is given by

$$\Delta \boldsymbol{z}_t = \mathbf{G} \cdot \boldsymbol{s}(\boldsymbol{z}_t), \tag{S9}$$

(eq. 3 of [10]) where $\mathbf{G}$ is the matrix of traits' genetic (co)variance (which is assumed to be constant during the evolution of the means so we can drop its time index for this section S2.2; in fact since we assume that each trait is encoded by a separate locus and all loci mutate in a similar way, it is reasonable to assume that $\mathbf{G} = \delta^2 \mathbf{I}$ here with $\mathbf{I}$ being the identity matrix), and

$$\boldsymbol{s}(\boldsymbol{z}_t) = \begin{pmatrix} s_a(\boldsymbol{z}_t) \\ s_b(\boldsymbol{z}_t) \\ s_d(\boldsymbol{z}_t) \\ s_h(\boldsymbol{z}_t) \end{pmatrix}, \tag{S10}$$

is the so-called selection gradient, which is a vector where each entry tells us whether selection favours an increase (when $s_u(\boldsymbol{z}) > 0$) or decrease (when $s_u(\boldsymbol{z}) < 0$) in the corresponding trait ($u \in \{a, b, d, h\}$) when the population average is $\boldsymbol{z}$. Such selection coefficient is given by

$$s_u(\boldsymbol{z}) = \frac{\partial w(\boldsymbol{z}_\bullet, \boldsymbol{z}_{-\bullet}, \boldsymbol{z})}{\partial u_\bullet} + (N-1) r_2 \frac{\partial w(\boldsymbol{z}_\bullet, \boldsymbol{z}_{-\bullet}, \boldsymbol{z})}{\partial u_2}, \tag{S11}$$

where here and hereafter, derivatives are evaluated where all individuals have the average phenotype, $\boldsymbol{z}_\bullet = \boldsymbol{z} = \boldsymbol{z}_2 = \ldots = \boldsymbol{z}$. The first term of eq. (S11) corresponds to the direct fitness effect of trait $u$: the marginal effect of a change in trait $u$ in the focal on its own fitness. The second term of eq. (S11), meanwhile, is the relatedness-weighted indirect fitness effect: the effect of a change in trait $u$ in a neighbour on the fitness of the focal individual, weighted by the coefficient $r_2$ of pairwise of relatedness, which is the probability that two individuals randomly sampled in a group are identical-by-descent, IBD, under neutrality. For the specific life-cycle described in the main text, this coefficient is given by

$$r_2 = \frac{1-m}{1+m(N-1)}, \tag{S12}$$

(e.g., [3] for the Moran model). As such, eq. (S11) can be seen as the marginal form of Hamilton's rule, $-c + r_2 b$, with direct effect as cost, $-c$, and indirect effect as a benefit, $b$.

**Singular strategy and convergence stability.** The dynamics given by eq. (S9) may eventually converge to an equilibrium, $\boldsymbol{z}^*$, so that the means no longer change (i.e. $\Delta \boldsymbol{z}_t = \mathbf{0}$). Such convergence first requires that

$$\boldsymbol{s}(\boldsymbol{z}^*) = \mathbf{0}. \tag{S13}$$

Indeed, because $\mathbf{G}$ is a positive-definite matrix (since it is a covariance matrix), condition eq. (S13) is the only way for $\boldsymbol{z}_{t+1} = \boldsymbol{z}_t = \boldsymbol{z}^*$. A strategy satisfying eq. (S13) is typically referred to as a singular strategy [13]. Whether a singular strategy is an attractor for directional selection (i.e. whether means will converge to $\boldsymbol{z}^*$ defined by eq. S13) can be investigated from the Jacobian matrix

$$\mathbf{J}(\boldsymbol{z}^*) = \begin{pmatrix} J_{aa}(\boldsymbol{z}^*) & J_{ab}(\boldsymbol{z}^*) & J_{ad}(\boldsymbol{z}^*) & J_{ah}(\boldsymbol{z}^*) \\ J_{ba}(\boldsymbol{z}^*) & J_{bb}(\boldsymbol{z}^*) & J_{bd}(\boldsymbol{z}^*) & J_{bh}(\boldsymbol{z}^*) \\ J_{da}(\boldsymbol{z}^*) & J_{db}(\boldsymbol{z}^*) & J_{dd}(\boldsymbol{z}^*) & J_{dh}(\boldsymbol{z}^*) \\ J_{ha}(\boldsymbol{z}^*) & J_{hb}(\boldsymbol{z}^*) & J_{hd}(\boldsymbol{z}^*) & J_{hh}(\boldsymbol{z}^*) \end{pmatrix} \tag{S14}$$

with $(u, v)$-entry

$$J_{uv}(\boldsymbol{z}^*) = \left.\frac{\partial s_u(\boldsymbol{z})}{\partial v}\right|_{z=z^*}. \tag{S15}$$

A *necessary* condition for a singular strategy to be an attractor is that the real parts of the eigenvalues of $\mathbf{J}(\boldsymbol{z}^*)$ are all negative. If so, we say that $\boldsymbol{z}^*$ is weakly convergence stable, "weakly" because it is still possible that the evolutionary dynamics do not converge towards $\boldsymbol{z}^*$ in the presence of genetic correlations among traits (i.e. there may exist non-diagonal matrix $\mathbf{G}$ so that iteration of eq. (S9) do not converge to $\boldsymbol{z}^*$). A *sufficient* condition for a singular strategy to be an attractor is that $\mathbf{J}(\boldsymbol{z}^*)$ is negative-definite [14, 15], i.e. that its symmetric part, $[\mathbf{J}(\boldsymbol{z}^*) + \mathbf{J}(\boldsymbol{z}^*)^{\mathrm{T}}]/2$ has only negative eigenvalues (where T denotes transpose; note that since $\mathbf{J}(\boldsymbol{z}^*) + \mathbf{J}(\boldsymbol{z}^*)^{\mathrm{T}}$ is symmetric with real entries, all its eigenvalues are real). A singular strategy $\boldsymbol{z}^*$ satisfying this sufficiency condition is said to be strongly convergence stable [15] in reference to the fact that as a result of mutation and selection, a population close to $\boldsymbol{z}^*$ will always gradually converge to $\boldsymbol{z}^*$, whatever the genetic correlations among traits (i.e, whatever the positive-definite $\mathbf{G}$ matrix). But since we assume that each trait is encoded by an independently mutating locus, there should be no genetic correlation among traits (under weak selection at least) . So the necessary condition for convergence stability should also be sufficient for trait averages to converge towards an equilibrium in our model.

#### S2.3 Stabilising, disruptive and correlational selection

**Hessian matrix.** Once the average traits in the population have converged to an equilibrium $\boldsymbol{z}^*$ under directional selection, the traits' (co)variances given by $\mathbf{G}$ around this population mean then start changing under the actions of mutations and selection. To the leading order of $\delta$, this change $\Delta\mathbf{G}_t = \mathbf{G}_{t+1} - \mathbf{G}_t$ over one demographic time step when the population mean is at a convergence stable phenotype is captured by

$$\Delta\mathbf{G}_t = \mathbf{M} + \mathbf{G}_t \cdot \mathbf{H}(\boldsymbol{z}^*) \cdot \mathbf{G}_t \tag{S16}$$

(eq. 3b of [10] with vanishing selection gradient), where the constant positive-definite matrix $\mathbf{M}$ captures the input of mutations (in the absence of pleiotropy and each trait mutating with the same probability and effects, as in our model, we can write this equation as $\mathbf{M} = \delta^2\mathbf{I}$), and the symmetric Hessian matrix

$$\mathbf{H}(\boldsymbol{z}^*) = \begin{pmatrix} H_{aa}(\boldsymbol{z}^*) & H_{ab}(\boldsymbol{z}^*) & H_{ad}(\boldsymbol{z}^*) & H_{ah}(\boldsymbol{z}^*) \\ H_{ba}(\boldsymbol{z}^*) & H_{bb}(\boldsymbol{z}^*) & H_{bd}(\boldsymbol{z}^*) & H_{bh}(\boldsymbol{z}^*) \\ H_{da}(\boldsymbol{z}^*) & H_{db}(\boldsymbol{z}^*) & H_{dd}(\boldsymbol{z}^*) & H_{dh}(\boldsymbol{z}^*) \\ H_{ha}(\boldsymbol{z}^*) & H_{hb}(\boldsymbol{z}^*) & H_{hd}(\boldsymbol{z}^*) & H_{hh}(\boldsymbol{z}^*), \end{pmatrix} \tag{S17}$$

captures the effects of selection. Each entry of this matrix informs on the nature of selection on traits' (co)variances at a singular strategy. Specifically, the sign of each diagonal entry indicates whether selection favours a decrease (when $H_{uu}(\boldsymbol{z}^*) < 0$) or increase (when $H_{uu}(\boldsymbol{z}^*) > 0$) in the variance of the corresponding trait (here $u$) when this trait evolves in isolation of the other traits [16]. In other words, $H_{uu}(\boldsymbol{z}^*)$ tells us whether selection on trait $u$ alone is stabilising (when $H_{uu}(\boldsymbol{z}^*) < 0$) or disruptive (when $H_{uu}(\boldsymbol{z}^*) > 0$). Further, the sign of each off-diagonal entry indicates whether selection favours a positive (when $H_{uv}(\boldsymbol{z}^*) > 0$) or negative (when $H_{uv}(\boldsymbol{z}^*) > 0$) correlation among the two corresponding traits (here $u$ and $v$) when these traits evolve in isolation from the others. The quantity $H_{uv}(\boldsymbol{z}^*)$ (with $u \neq v$) has accordingly been coined as the coefficient of "correlational selection" [16].

Under our life-cycle assumptions (i.e. Moran reproductive process), the $(u, v)$-entry of the Hessian matrix is given by

$$
\begin{aligned}
H_{uv}(\boldsymbol{z}^*) = \frac{\partial^2 w(\boldsymbol{z}_\bullet, \boldsymbol{z}_{-\bullet}, \boldsymbol{z})}{\partial u_\bullet \partial v_\bullet} &+ (N-1) r_2 \left[ \frac{\partial^2 w(\boldsymbol{z}_\bullet, \boldsymbol{z}_{-\bullet}, \boldsymbol{z})}{\partial u_2 \partial v_2} + \frac{\partial^2 w(\boldsymbol{z}_\bullet, \boldsymbol{z}_{-\bullet}, \boldsymbol{z})}{\partial u_\bullet \partial v_2} + \frac{\partial^2 w(\boldsymbol{z}_\bullet, \boldsymbol{z}_{-\bullet}, \boldsymbol{z})}{\partial u_2 \partial v_\bullet} \right] \\
&+ (N-1)(N-2) r_3 \frac{\partial^2 w(\boldsymbol{z}_\bullet, \boldsymbol{z}_{-\bullet}, \boldsymbol{z})}{\partial u_2 \partial v_3}
\end{aligned}
\tag{S18}
$$

(eq. 7.b of [10]), where derivatives are all evaluated at the singular phenotype, $\boldsymbol{z}_\bullet = \boldsymbol{z} = \boldsymbol{z}_2 = \ldots = \boldsymbol{z}^*$, and

$$
r_3 = \frac{2(1-m)}{2+m(N-2)} r_2 = \frac{2(1-m)^2}{[1+m(N-1)][2+m(N-2)]}
\tag{S19}
$$

is the three way relatedness coefficient for the Moran model, i.e., the probability that three individuals randomly sampled from the same group are IBD under neutrality (more generally, the Hessian matrix eq. S17 is comprised of extra terms that capture the effects of traits on relatedness, eq. 7.a & c of [10], but these vanish under fecundity effects under a Moran life-cycle at a singular phenotype, eq. 16 of [3] for details, and so we can ignore them here).

**Stabilising and disruptive selection.** With all traits coevolving, whether selection is: (1) stabilising, keeping traits' (co)variances small so that the phenotypic distribution remains unimodal around the equilibrium; or (2) disruptive, favoring an increase in the variance of some traits (and possibly some covariances), depends on the leading eigenvalue of the Hessian matrix $\rho(\mathbf{H}(\boldsymbol{z}^*))$ (where $\rho(\mathbf{A})$ denotes the leading eigenvalue of a matrix $\mathbf{A}$). Selection is stabilising when $\rho(\mathbf{H}(\boldsymbol{z}^*)) < 0$. In this case, selection purges genetic variation that deviates from the singular strategy. Such a strategy $\boldsymbol{z}^*$ is said to be uninvadable. As a result of stabilising selection combined with mutation, traits in the population reach an equilibrium that is characterised by a distribution concentrated around the singular strategy. By contrast, selection is disruptive when $\rho(\mathbf{H}(\boldsymbol{z}^*)) > 0$. In this case, genetic variation increases along the eigenvector associated with the leading eigenvalue. This may lead to evolutionary branching whereby the phenotypic distribution goes from being unimodal to bimodal so that two highly differentiated morphs or types coexist in the population (for further considerations on this when multiple traits coevolve, [17]).

The analysis of the eigenvalue $\rho(\mathbf{H}(\boldsymbol{z}^*))$ can be prohibitively complicated. Fortunately there exists simpler conditions that are sufficient for disruptive selection to occur (i.e. for $\rho(\mathbf{H}(\boldsymbol{z}^*)) > 0$), which use the fact that the Hessian is a symmetric matrix [18]. In particular, if any diagonal entry is positive ($H_{uu}(\boldsymbol{z}^*) > 0$), then $\rho(\mathbf{H}(\boldsymbol{z}^*)) > 0$, i.e. if selection is disruptive on any trait when it evolves in isolation from the others, then selection is disruptive when they all co-evolve. Alternatively when $H_{uu}(\boldsymbol{z}^*) < 0$ for all $u$, selection is disruptive ($\rho(\mathbf{H}(\boldsymbol{z}^*)) > 0$) if the off-diagonal entry of any $2\times2$ submatrix of $\mathbf{H}(\boldsymbol{z}^*)$ is large relative to the diagonal entries of this submatrix so that $H_{uv}(\boldsymbol{z}^*)^2 > H_{uu}(\boldsymbol{z}^*)H_{vv}(\boldsymbol{z}^*)$ (for some $u \neq v$). Put differently, selection is disruptive if correlational selection among two traits is large relative to stabilising selection on both isolated traits.

### S3 Analyses

In this supplement, we detail the mathematical analyses underlying the results summarized in the main text. In particular, we derive the singular strategies (eq. 9 of the main text), show that we expect these strategies to be convergence stable but not uninvadable, i.e., that we expect evolutionary branching to happen in our model. The basis of all our results is obtained by first computing payoff (substitute eqs. (1)–(3) into eq. (S1)) that is substituted into eq. (S5) to calculate fitness, which is in turn substituted into the selection gradient vector eq. (S11) and Hessian matrix eq. (S18). All the relevant quantities for our evolutionary analysis unfold from these operations. We provide a Mathematica notebook to follow and check all computations reported below ([19], see attached M-file).

#### S3.1 Helping and belligerence

To begin with, we derive the conditions under which belligerence emerge (eq. 6 of main text). First, we set the selection gradient on helping to zero when belligerence and both forms of bravery are absent in the population, i.e. we set $s_h(\boldsymbol{z}) = 0$ with $\boldsymbol{z} = (0, 0, 0, h)$. After rearrangements, we obtain eq. (6) of the main text which gives the first order condition to the equilibrium of helping when the other traits are absent. Condition eq. (6) shows that helping equilibrium increases with the parameter $\kappa^{\mathrm{R}}$ which increases with relatedness (eq. 7, Fig. 4). This parameter $\kappa^{\mathrm{R}}$ in fact incorporates two antagonistic effects of limited dispersal on the evolution of helping or other pro-social traits. On one hand, high relatedness due to limited dispersal favors prosocial behavior within groups because in this case, the recipients of the actions of an individual tend to bear the same genes underlying those actions (i.e. kin selection operates). On the other hand, group members typically also compete more strongly for the same local resources than two randomly sampled individuals in the populations (which is the case in our model, section 2.1). As a result, the positive effects of relatedness on the evolution of behaviour tend to be mitigated by competition between relatives, referred to as "kin competition". This is typically reflected in models of social evolution under limited dispersal where evolutionary stable trait values depend on relatedness scaled by local competition, which is captured by the parameter $\kappa < r_2$ (eq. 7, Fig. 4; e.g., [2, 20–24]). As mentioned in the main text, our condition for helping evolution depends on $\kappa^{\mathrm{R}}$ rather than $\kappa$ because the benefits of helping are shared equally within the group. An individual therefore always recoup a share $1/N$ of its own investment, which increases selection on helping (i.e. $\kappa^{\mathrm{R}} > \kappa$, eq. 7, Fig. 4). Since all traits we study are in effect pro-social and benefit the whole group equally, this quantity $\kappa^{\mathrm{R}}$ will also emerge in the selection gradients of the other traits other than helping ($a$, $b$ and $d$).

Second, we look at where the selection gradient on belligerence is positive when helping is present in the population but bravery is not, i.e. look at where $s_a(\boldsymbol{z}) > 0$ with $\boldsymbol{z} = (0, 0, 0, h)$. This gives us eq. (8) of the main text.

#### S3.2 All traits co-evolving

##### S3.2.1 Singular strategies

Recall that in order to go further in our analysis and analyse the case where all traits are co-evolving, we make the assumptions: (1) $B(Nh) = \beta\sqrt{Nh}$ (where $\beta > 0$ is a constant); (2) $\alpha(a) = a$; (3) $g(b) = b$; (4) $C(h, b, d) = h + c_{\mathrm{b}} b + c_{\mathrm{d}} d$; and (5) $F(\pi) = \pi$.

First, the singular values for offensive and defensive bravery (eqs. 9b-9c) are found by setting the selection gradients for offensive and defensive bravery to zero and solving these equations for $b^*$ and $d^*$ (i.e. find $b^*$ and $d^*$ in $\boldsymbol{z}^* = (a^*, b^*, d^*, h^*)$ such that $s_b(\boldsymbol{z}^*) = s_d(\boldsymbol{z}^*) = 0$). The singular value for helping (eq. 9d) is in turn found by solving $s_h(\boldsymbol{z}^*) = 0$ for $h^*$ where $b^*$ and $d^*$ are given by eqs. (9b)-(9c). Similarly, the (implicit) singular value for belligerence (eq. 9a) is found by solving $s_a(\boldsymbol{z}^*) = 0$ for $a^*$ where $b^*$ and $d^*$ are given by eqs. (9b)-(9c).

##### S3.2.2 Connections with previous results on bravery evolution

The equilibrium for bravery in our model (eq. 9b-9c) is consistent with previous models. In particular, when fighting is certain (setting $\phi = 1 - e^{-a^*} = 1$), offensive and defensive bravery costs are equal ($c_\mathrm{b} = c_\mathrm{b} = c$), and there is no inherent advantage to being in an offensive or defensive position ($\omega = 1/2$), eqs. (9b)-(9c) reduce to

$$b^* = d^* = \frac{1}{4} \times \frac{B}{Nc} \times \kappa^\mathrm{R}. \tag{S20}$$

Under complete dispersal (i.e. random group formation, $m = 1$), we have $\kappa^\mathrm{R} = 1/N$ (eq. 7) so that eq. (S20) reads as, $b^* = d^* = (1/4)B/(cN^2)$. This is equal to the equilibrium found in classical models of investment into contest (e.g. eq. 10 of [25], first equation p. 1018 of [26]). Such congruence follows from the connection between our payoff function and that used in classical model (eq. 5).

Rusch and Gavrilets [26] also present an expression for investment into contest where it is claimed groups consists of relatives due limited dispersal (their first equation p. 1023 of ref. [26], referred to as eq. (RG) hereafter). This eq. (RG) is inconsistent with our eq. (S20). While the exact source of this inconsistency is not fully clear to us, there are several problems with eq. (RG). First, it is in conflict with [26]'s own equation without relatedness (i.e. in the same paper, on p. 1018), as eq. (RG) with $r_2 = 0$ does not reduce to the latter. Second, it disagrees with the notion that when $r_2 = 1$, the equilibrium strategy should maximizes group payoff (as there is no conflict within groups of clones). Third, eq. (RG) was taken from [27] in which there are several discrepancies between biological assumptions and fitness accounting[1]. These issues lead us to believe that eq. (RG) is erroneous.

##### S3.2.3 Explicit solutions

Our implicit expressions for the equilibria (eq. 9) highlight the inter-dependence between the four co-evolving traits. To obtain explicit expressions in terms of model parameters only, we substitute for $h^*$ (eq. 9d) into eq. (9a) and solve the resulting equation yielding

$$a^* = \log\left(\frac{c_2 - 2c_1 + {v^*}^2\beta^2\kappa^\mathrm{R}/2}{c_2 - c_1 - v^*(1 - v^*)\beta^2\kappa^\mathrm{R}/2}\right). \tag{S21}$$

Substituting eq. (S21) into eq. (9d) and in turn these into eqs. (9b)-(9c) gives explicit solutions for the equilibria of the other traits.

[1] For e.g.: fitness in the model of [27] goes to zero when the number of groups in the population becomes large, see their eqs. (2) and (4); the "correction" factor in their eq. 6, i.e. their eq. 13, is just stated, it is neither derived nor supported by reference to previous literature; the equation for relatedness (above their eq. 14) is for a model with isolation by distance but nowhere in the manuscript is such isolation by distance evoked and the equations are more consistent with uniform dispersal.

##### S3.2.4 Convergence stability

From our general assumption that benefits of the common good decelerate ($B''(h) < 0$) but that costs associated with obtaining this common good (either through production or attacking) do not, it seems reasonable to expect that traits will not grow indefinitely. In other words, we expect that provided helping and belligerence emerge (section 3.1), the joint singular trait value given by eqs. (9) is an evolutionary attractor (i.e. convergence stable). Although the Jacobian matrix (eq. S14) for our model is too complicated to check this expectation analytically, a numerical approach supports it (Mathematica Notebook for results). Indeed, when we sampled $10^6$ random combinations of model parameters such that helping and belligerence emerged, we found that in 99.9% of cases the real part of the dominant eigenvalue of the Jacobian matrix at the singular value was negative (i.e. that $\boldsymbol{z}^*$ is – weakly – convergence stable), and that for 55% of those combinations, the Jacobian matrix was further negative-definite (i.e. that $\boldsymbol{z}^*$ is – strongly – convergence stable). This tells us that in the majority of cases, the population will gradually converge to a state where its phenotypic mean is given by the singular strategy $\boldsymbol{z}^*$, whatever the genetic correlations among traits. If traits are not genetically correlated, which should be the case under our assumption that each trait mutates independently (provided selection is not too strong), such convergence should happen in essentially all cases.

##### S3.2.5 Local evolutionary stability

The above analysis suggests that first, the mean phenotype in the population will converge to the singular value $\boldsymbol{z}^*$ while the traits' (co)variances remain small. Selection on these (co)variances then depend on the Hessian matrix $\mathbf{H}(\boldsymbol{z}^*)$ (eqs. S17–S18). Although the Hessian matrix have complicated entries for our model, it turns out to have a simple sign structure,

$$\mathbf{H}(\boldsymbol{z}^*) = \begin{pmatrix} 0 & >0 & 0 & 0 \\ >0 & <0 & 0 & 0 \\ 0 & 0 & <0 & >0 \\ 0 & 0 & >0 & <0 \end{pmatrix} \tag{S22}$$

(Mathematica Notebook). This sign structure tells us a few things. The first is that since none of the diagonal element is positive, none of the traits are under disruptive selection when they evolve in isolation from one another. Further, since $H_{aa}(\boldsymbol{z}^*) = 0$, selection on bravery alone is neither stabilizing nor disruptive at the singular strategy, and this holds for all scenarios investigated in this paper (section S5). This entails that in the $2\times2$ upper left submatrix, $H_{ab}(\boldsymbol{z}^*)^2 > H_{aa}(\boldsymbol{z}^*)H_{bb}(\boldsymbol{z}^*) = 0$, which means that whenever belligerence coevolves with offensive bravery, selection is disruptive favouring polymorphism. In addition, since $H_{ab}(\boldsymbol{z}^*) > 0$ and $H_{dh}(\boldsymbol{z}^*) > 0$, we expect this polymorphism to be characterised by a positive correlation between belligerence and offensive bravery, and between helping and defensive bravery. Note that because the Hessian matrix provides information on the nature of selection locally, i.e. based on the assumption that the phenotypic distribution is peaked around the singular strategy, our conclusions on correlations hold at least for when the polymorphism initially emerges. As a result, we cannot say anything about the long term nature of the polymorphism [17]. Our simulations provide nonetheless insights into this (Figs. 2-3).

The key feature of eq. (S22) that promotes the emergence of polymorphism is the fact that $H_{aa}(\boldsymbol{z}^*) = 0$. This property will typically not hold when warfare is modelled as a reproductive strategy. This can be seen by considering that when eq. (S6) is substituted into eq. (S18), we are likely to have $H_{aa}(\boldsymbol{z}^*) < 0$. In other words, selection on belligerence alone will tend to be stabilising and thus inhibit disruptive selection.

### S4 Individual based simulations

To confirm our mathematical analysis and investigate trait associations in the longer term, we used individual based stochastic simulations (with finite number of groups $N_g < \infty$). Such simulations have been carried out extensively across several papers, in which they been shown to generally be in excellent agreement with results from local analyses in group-structured populations, irrespective of group-size and dispersal as long as $m > 0$ and $N_g$ is sufficiently large for genetic drift to be negligible ([3, 10, 28, 29]; for discussions on the effects of drift on polymorphism in well-mixed populations, see [30, 31]). With this in mind and given the time taken for such simulations to run, we focused simulations here on a specific set of parameter values representative of our model.

Our individual based simulations follow a population composed of $N_g = 1250$ groups, each populated by $N = 8$ individuals, using Mathematica 10.2.0.0 (see attached M-file, [19]). Starting with a monomorphic population, we track the evolution of the phenotypic distribution for a fixed number of generations. Each individual $i \in \{1, \ldots, N_g N\}$ at each generation is characterised by a vector of traits $(a_i, b_i, d_i, h_i)$. At the beginning of a generation, we first calculate the payoff $\pi_i$ of each individual according to its traits, those of its neighbours and the average traits in the population (using eqs. S1-S3). Fecundity is taken as payoff (i.e. $F(\pi_i) = \pi_i$). We also ran simulations where we explicitly modelled individual battles following the (finite) island model of warfare [6] so that individual fecundity depended on a specific sequence of events (e.g. whether a raid took place, whether it was won, how many units of resources were present in the specific group raided). As expected from the considerations of section S1.2, these simulations were consistent with those where fecundity was given by expected payoff (using eqs. S1-S3) provided baseline fecundity $v_b$ was high enough. Since the former were more stochastic and significantly more time consuming, we focused on the latter (i.e. using eqs. S1-S3 to calculate fecundity).

After fecundity is calculated, an individual is randomly sampled in each group to be replaced by an offspring. This offspring is then chosen independently in each group by sampling among the population an individual according to its group and the group in which an breeding spot is being filled. Specifically, if an individual belongs to the same group in which the breeding spot is filled, then its weight is $\pi_i(1-m)$, where $m$ is the dispersal probability. If it belongs to another group, then its weight is $\pi_i m/(N_g - 1)$. Once an individual is chosen to fill the breeding spot, each of its traits mutate independently with probability $\mu = 0.01$. If a trait does not mutate, then it has the same value as in the parent. If a trait does mutate, then we add to parental values a small perturbation that is sampled from a normal distribution with mean 0 and variance $\sigma^2 = 0.02^2$. The resulting phenotypic values are truncated to remain positive (and less than 1 for belligerence $a$ as it is a probability in our examples). We repeat the procedure for a fixed number of generations (Figure legends for parameter values used).

### S5 Robustness of results

In addition to exploring different individual cost function (eq. 11 in main text), we relaxed our baseline model (detailed at the beginning of section 3.2) in three other directions, and (1) considered two further contest functions that have been suggested in the literature: $g(y) = y^\lambda$ and $g(y) = \exp(\lambda y)$ (e.g., [32, 33]); (2) explored the effect of group decision by modeling it as majority "voting" $\alpha(a_0) = a_0^\lambda / [a_0^\lambda + (1-a_0)^\lambda]$, so that as $\lambda$ increases, the decision to raid increasingly becomes binary according to whether the group-average of belligerence $a_0$

is below or above 1/2; and finally (3) allowed for a sigmoidal relationship between total investment into helping in the group and the benefits of this common-pool resource, with $B(Nh) = \beta(Nh)^{\lambda}/((Nh)^{\lambda} + \chi)$. We find that in all examined cases, these different functions influence the value of the equilibrium for each trait in a quantitative way (Suppl. Fig. 4A-D) but not the qualitative nature of these equilibria. Those are still internal evolutionary attractors under directional selection when the costs of fighting are non-additive ($c_2 > 2c_1$), and once the population has converged to the joint equilibrium, selection becomes disruptive favouring the emergence of polymorphism (Suppl. Fig. 5A-D). Our analysis additionally indicates that this polymorphism should again be characterised by a positive association between belligerence and offensive bravery (as indicated by a positive correlational selection coefficient among these two traits, Suppl. Fig. 5A-D).

These extensions suggest that the polymorphism we observed under our baseline model assumptions (section 3.2) is robust to changing the behavioral rules of within- and between-groups interactions. Variations of the demographic assumptions are also unlikely to change these results. We chose the Moran process for simplicity but of course many different alternative life-cycle assumptions are possible (e.g., all individuals die per time step, each individual survives with a fixed probability; density-dependent regulation occurs before dispersal "soft-selection"; dispersal occurs through propagules of individuals). Yet we know from previous social evolution models that these alternatives do not qualitatively affect equilibrium conditions (eq. 9), as all these life-cycle variations can be accounted by varying the scaled-relatedness coefficient [23, 24]. Such variations are also unlikely to qualitatively alter the analysis of disruptive selection (and therefore polymorphism) as disruptive selection can also be expressed in terms of summary demographic variables [3].

One particularly strong assumption when applying our model to animals is that individuals are haploid and reproduce asexually. Thankfully, neither the condition for equilibrium nor for disruptive selection will be qualitatively influenced by diploidy and sexual reproduction when genes have additive effects within individuals [2, 34]. The emergence of polymorphism due to correlational selection may however depend on the genetic architecture of traits [29]. Nonetheless, if the genetic architecture of belligerence and bravery are such that their associations are heritable (e.g. tightly linked or encoded by the same pleiotropic locus), or alternatively if such architecture is allowed to evolve, then the emergence of polymorphism will unfold as in our model [29, 34–36].

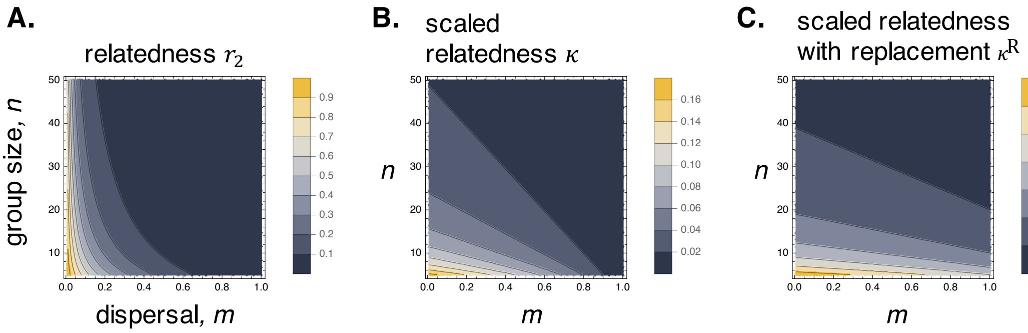


**Supplementary Figure 1: (Scaled) Relatedness, with and without replacement.** A. Relatedness $r_2$ (eq. S12), in the island model when a single reproductive spot is replaced in each generation; B. Scaled relatedness $\kappa$ (eq. 7); C. Scaled relatedness with replacement $\kappa^R$ (eq. 7) as a function of dispersal $m$ and group size $n$ (legend for values).

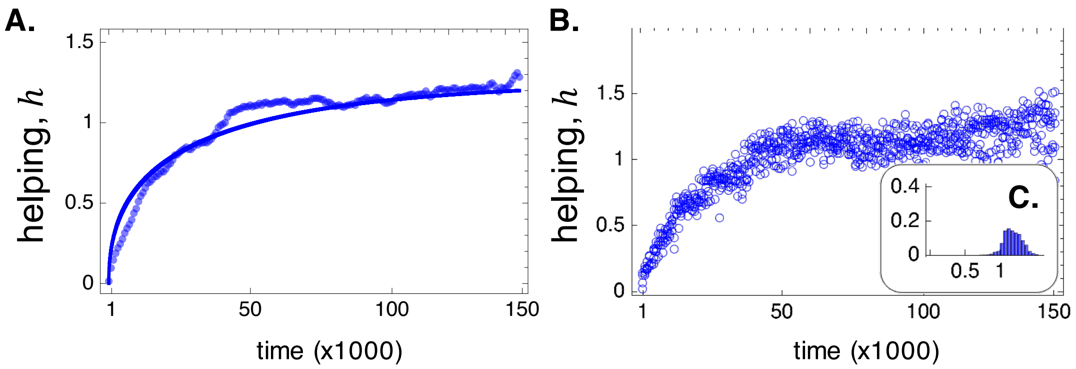


**Supplementary Figure 2: The evolution of helping.** **A**: The average level of helping as a function of demographic time $t$ (dots: observed in simulations, section S4 for details; full line: analytical predictions from eq. (S9) with variance-covariance matrix $\mathbf{G}$ composed of all zeroes except $G_{hh} = 0.00375$, chosen heuristically) when the other traits are absent in the population, i.e., $a = d = b = 0$ for all individuals throughout; with $B(Nh) = \beta\sqrt{Nh}$, $C(h,b,d) = h$, $\beta\sqrt{N} = 100$, $\upsilon_{\rm b} = 0$, $N = 8$, $m = 0.476$. **B**: Individual values of helping observed in a simulation (shown for 5 individuals randomly sampled every 800 time points). **C**: Distribution of helping in a simulated population (calculated from time 100'000 for 50'000 time steps).

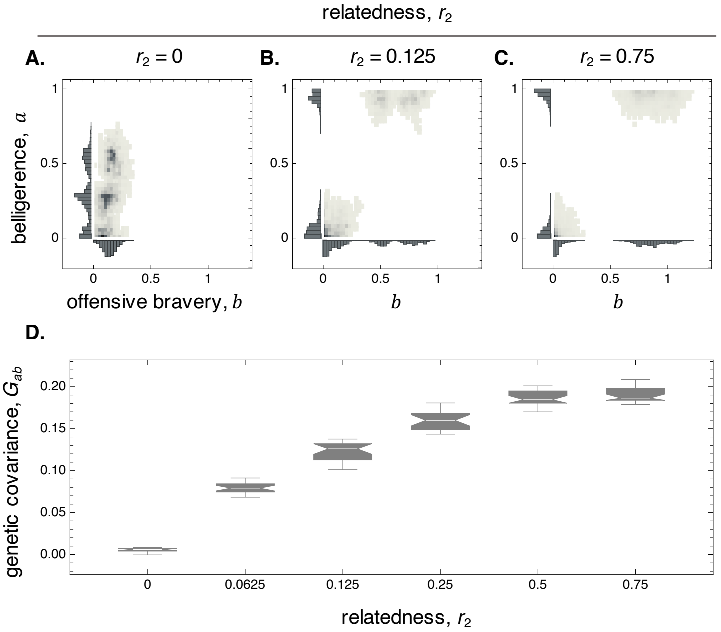


**Supplementary Figure 3: The effect of limited dispersal and relatedness on polymorphism.** **A-C** Joint distribution of belligerence and offensive bravery in simulated populations at equilibrium for different levels of relatedness (found by fixing $N = 8$ and varying $m$ in $r_2$ eq. (S12)) with: A. $r_2 = 0$ (so $m = 1$); B. $r_2 = 0.125$ (so $m = 0.467$); $r_2 = 0.75$ (so $m = 0.04$) (other parameters, same as Fig. 1 middle; joint distribution calculated over 100'000 time points after 150'000 of evolution). **D.** Distribution of covariance between belligerence and offensive bravery over time at equilibrium according to relatedness within groups (same as A other than $m$).

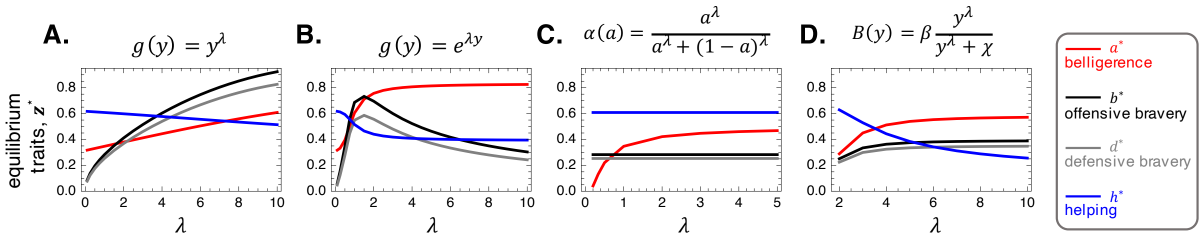


**Supplementary Figure 4: The effect of changing functional relationships on equilibrium.** Equilibrium value of each trait ($a$ in red, $b$ in black, $d$ in gray, $h$ in blue) against $\lambda$ that is used as the parameter in the different following function: **A.** $g(y) = y^{\lambda}$; **B.** $g(y) = \exp(\lambda y)$; **C.** $\alpha(a_0) = a_0^{\lambda}/[a_0^{\lambda} + (1-a_0)^{\lambda}]$; **D.** $B(Nh) = \beta(Nh)^{\lambda}/((Nh)^{\lambda} + \chi)$ (with $\beta = 100$ and $\chi = 10$, varying $\lambda$ for the steepness of the sigmoid; unless otherwise stated: $\alpha(a) = a$, $g(y) = y$, and $B(Nh) = \beta\sqrt{Nh}$; Other parameters: $C(h,b,d) = h^2 + c_{\mathrm{b}}b^2 + c_{\mathrm{d}}d^2$, $c_1 = 18$, $c_2 = 115$, $c_{\mathrm{d}} = 1$, $c_{\mathrm{b}} = 0.8$, $\omega = 0.5$, $\beta\sqrt{N} = 100$, $v_{\mathrm{b}} = 0$, $N = 8$, $m = 0.467$ so that $r_2 = 0.125$). All computed from solving the selection gradients numerically for singular values (see Mathematica Notebook). All these equilibria strategies are at least weakly convergence stable (see text below eq. S14) but not locally stable due to correlational selection among belligerence and offensive bravery (Suppl. Fig. 5). This indicates that they are attractors under directional selection but that once the population expresses these traits on average, selection becomes disruptive, favoring an association between belligerence and offensive bravery.

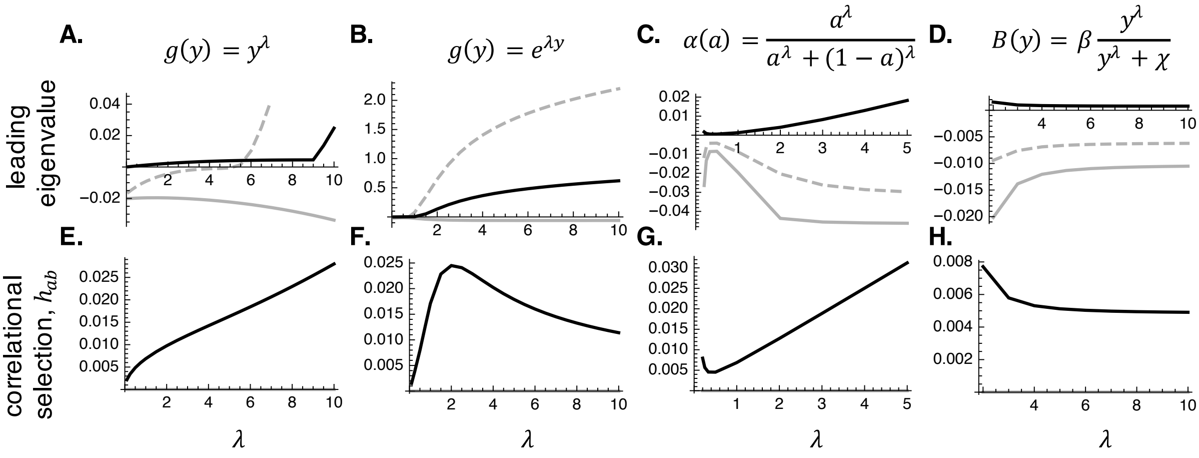


**Supplementary Figure 5: Summary analysis of the effect of contest types and group decisions. A-D** Leading eigenvalues of the Jacobian (in dashed gray), symmetric part of the Jacobian (in full gray), and Hessian (in black), showing that the singular values plotted in Fig. 4 are all (at least weakly) convergence stable but none is locally stable. This suggests that polymorphism also emerges under these different assumptions (Fig. 4 for details on parameters values). **E-H** Correlational selection among belligerence and offensive bravery at the singular strategy. It is always positive, highlighting that selection still favours a positive selection.
