## Supplementary material S1-S5, figures, matehmatica notebook and code. for "Evolution of warfare by resource raiding favors polymorphism in belligerence and bravery": Individual_based_simulations.m

```
(* This m-file runs individual based simulations used in the manuscript entitled "On the evolution of warfare by resource raiding and the emergence of polymorphism in belligerence and bravery" by Charles Mullon and Laurent Lehmann. Overall simulation procedure is described in the Electronic Supplementary Material S4. They were run using Mathematica v.12.1.1.0. To run on one's own machine, make sure to adapt the output files to your own directories via the variables streamPop and streamMeans below. *)

Clear["Global`*"];
 
 (* The function "iteration1Gen" defined below takes for input "populationPhenotypesWithGeneration" which is a list whose first element is a generation mumber, say t, and the second the whole list of traits in the population at this generation, L(t). Its output is then a list whose first element is the next generation mumber, t+1, and the second L(t+1). We will iterate this function for a fixed number of generations for given parameters and initial population L(0). *)

iteration1Gen[populationPhenotypesWithGeneration_] :=
Block[
(* define local variables *)
{generationN = populationPhenotypesWithGeneration[[1]],populationPhenotypes = populationPhenotypesWithGeneration[[2]],MeanDemesx,MeanDemesy,Fertility,parapatricFitness,x1,y1,x,y,event,PhilopatricFitness,fitness,Meanx,Meany,Varx,Vary,Covarxy,pop,mutation,Mutationcounter = 0,MutationMatrix,NewGeneration,pos},
      
      (* generation *)
      PrintTemporary[generationN];
      generationN++;
      
      (* every "sampling" generations, print the phenotypes of all individuals in the population in file "streamPop" *)
      If[Divisible[generationN, sampling],
         PutAppend[populationPhenotypes, streamPop]
         ,];
      
      (* Calculate mean phenotype in each group *)
      (* belligerence *)
      a0R = Map[Mean, populationPhenotypes[[All, All, 1]]];
      (* offensive bravery *)
      b0R = Map[Mean, populationPhenotypes[[All, All, 2]]];
      (* defensive bravery *)
      d0R = Map[Mean, populationPhenotypes[[All, All, 3]]];
      (* helping *)
      h0R = Map[Mean, populationPhenotypes[[All, All, 4]]];
      
      (* Calculate mean traits in whole pop *)
      (* belligerence *)
      a1 = Mean[ a0R ];
      (* offensive bravery *)
      b1 = Mean[ b0R ];
      (* defensive bravery *)
      d1 = Mean[ d0R ];
      (* helping *)
      d1 = Mean[ h0R ];
      
      (* print the mean phenotype every generation in file "streamMeans" *)
      PutAppend[{a1,b1,d1,d1}, streamMeans];
      PrintTemporary[{a1,b1,d1,d1} ] ;
    
      (* Calculate payoff/fertility of each individual in the population with function f which is defined below according to eqs.(S1-1)-(S1-3) from Electronic Supplementary Material S1 *)
      Fertility =  Table[ f[ populationPhenotypes[[i,j]], { a0R[[i]], b0R[[i]] ,d0R[[i]] , h0R[[i]]  } ,{a1,b1,d1,d1} ], {i, Nd}, {j, Nind}];
      
      (* calculate parapatric fitness, i.e. number of offpsring sent by each individual in a random patch *)
      parapatricFitness = m*Fertility/(Nd-1) ;
      
      (* determine the single adult that dies in each group and that will be replaced by an offspring  *)
      deads = RandomInteger[{1, Nind}, Nd];
      
      (* Determine  the successful offspring in each group i *)
      NewGeneration = Table[
                            (* calculate philopatric fitness for adults belonging to group i  *)
                            PhilopatricFitness = Fertility[[i]]*(1-m);
                            (* determine the fitness of each adult in the population with respect to group i *)
                            fitness =  ReplacePart[ parapatricFitness, i -> PhilopatricFitness]  ;
                            (* Replace the dead adult by a randomly chosen offspring where sampling is weighted by fitness *)
                             Table[ If[ deads[[ i ]] == j ,   RandomChoice[ Flatten[ fitness ] -> Flatten[populationPhenotypes, 1]] , populationPhenotypes[[i,j]] ], {j, Nind} ]
                                , {i, Nd}
                                    ];
      
      (* compute the effect of mutation on each succesful offpsring via the matrix "MutationMatrix"  *)
      event = RandomChoice[{1 - Pmutation, Pmutation} -> {0, 1}, {Nd,4}];
      mutation = RandomVariate[NormalDistribution[0, mutationStep ],Total[Flatten[event]]];
      MutationMatrix = Table[ If[ deads[[ i ]] == j ,
                                 Table[
                                       If[ event[[i,trait]] == 0 , 0 ,
                                          Mutationcounter++;
                                          mutation[[Mutationcounter]]
                                          ] ,{trait, 4} ]
                                 ,{0,0,0,0} ]
                             , {i, Nd} , {j, Nind}
                             ];
      
      (* Add the effects of mutation on population phenotypes *)
      NewGeneration = NewGeneration + MutationMatrix ;
      (* truncate the population phenotypes so that belligerence is between 0 and 1 and all the other traits are positive *)
      NewGeneration = MapAt[ Clip[#, {minT, 1.}] &, NewGeneration , {All, All, 1}];
      NewGeneration = MapAt[ Clip[#, {minT, Infinity}] &, NewGeneration , {All, All, 2}];
      NewGeneration = MapAt[ Clip[#, {minT, Infinity}] &, NewGeneration , {All, All, 3}];
      NewGeneration = MapAt[ Clip[#, {minT, Infinity}] &, NewGeneration , {All, All, 4}];
      
      (* output *)
      Prepend[{NewGeneration}, generationN]
      ];

(* Demographic parameters *)
Nind = 8 ; (* number of individual per groups *)
m = 0.466667 ; (* dispersal, so with Nind = 8 and m = 0.466667, pariwise relatedness r2 = 0.125 *)
Nd = 125 ; (* number of groups -- 1250 in our manuscript *)
Ngenerations = 100 ; (* Total number of generations to simulate -- up to 300'000 in our manuscript *)
sampling = 100*Nind ; (* sample whole population every ~ 100 generations *)

(* Costs and benefits *)
vb = 0. ; (* Baseline payoff *)
beta = 100. ; (* Benefit factor of common good *)
c1 = 18. ; (* Cost of one contest *)
c2 = 115.  ; (* Cost of two contests *)
cb = 0.51 ; (* Individual cost of offensive bravery *)
cd = 0.67 ; (* Individual cost of defensive bravery *)
om = 0.5 ; (* weight of offense vs defense in contest *)

(* contest function *)
g[y_]:= y;
                      
(* Cost function *)
Cc[bi_,di_,hi_] := hi + cb bi + cd di ;
                      
(* Helping benefits *)
Bb[nh_] := beta Sqrt[nh] ;
                      
(* minimum threshold to avoid divisions by zero *)
minT = 10^(-6.) ;
                      
(* fecundity function of a focal individual with traits {ai,bi,di,hi} in a group with average traits {a0,b0,d0,h0} in a population with average {a1,b1,d1,h1} according to eqs.(S1-1)-(S1-3) from Electronic Supplementary Material S1, here for alpha(a) = a *)
f[{ai_,bi_,di_,hi_},{a0_,b0_,d0_,h0_},{a1_,b1_,d1_,h1_}] :=  Max[ minT , vb + (c1 (-1+E^-a1)+(E^-a1 Bb[h0 Nind] (om g[b1 Nind]-E^a1 (-1+om) g[d0 Nind]))/(om g[b1 Nind]-(-1+om) g[d0 Nind])+(a0 E^(-2 a1) (-1+E^a1) (om (c2-c2 E^a1+c1 (-2+E^a1)+E^a1 Bb[h1 Nind]) g[b0 Nind]-(c2-c2 E^a1+c1 (-2+E^a1)) (-1+om) g[d1 Nind]))/(a1 (om g[b0 Nind]-(-1+om) g[d1 Nind])))/Nind - Cc[bi,di,hi]  ];


(* Mutation parameters *)
Pmutation = 0.01 ;  (* probability of a mutation per locus *)
mutationStep = 0.02;  (* standard deviation in phenotypic effect of mutation *)

(* Initial population *)
aInit = minT ; (* inital belligerence *)
bInit = minT ;  (* inital offensive bravery *)
dInit = minT ; (* inital defensive bravery *)
hInit = minT ; (* inital helping *)

(* inital input *)
InitialPopulation = Prepend[{ Table[{aInit, bInit, dInit , hInit} , {i, Nd}, {j, Nind}]  }, 0] ;

(* Output files *)
   streamPop = OpenWrite["Pop_IBS.csv", PageWidth -> Infinity];
   streamMeans = OpenWrite["Means_IBS.csv", PageWidth -> Infinity];
                      
(* launch simulations *)
   Nest[iteration1Gen, InitialPopulation, Ngenerations];

(* Close files and end program *)
   Close[streamPop];
   Close[streamMeans];

Quit[];
```
